# Antibody feedback establishes an affinity brake in the germinal center

**DOI:** 10.1101/2025.11.13.688298

**Authors:** Yu Yan, Xuesong Wang, Zhenfei Xie, Daniel L. V. Bader, Ryan H. Lim, Krystal M. Ma, Christopher A. Cottrell, Jon M. Steichen, Liling Xu, Paula Maldonado Villavicencio, Madhav Akauliya, Ja-Hyun Koo, Jacqueline Ming Shen, Alexandra Vernich, Oleksandr Kalyuzhniy, Joel D. Allen, Ali A. Albowaidey, Anthony Alicea, Bingxian Chen, Erik Georgeson, Jordan Renae Ellis-Pugh, Nushin Alavi, Abigail Esposito, Hannah Naili, Nicole Phelps, Brendon Kelley, Michael Kubitz, Quynh Anh Phan, Alessia Liguori, Thavaleak Prum, Ryan Tingle, Danny Lu, Saman Eskandarzadeh, Xiaotie Liu, John E. Warner, Stephanie R. Weldon, Sunny Himansu, Max Crispin, Usha Nair, Sophia Liu, William R. Schief, Facundo D. Batista

## Abstract

Recent advances in mRNA vaccine technology have opened the door to novel types of antigen display, but little is yet known as to how B cells recognize and respond to these formats. By delivering an mRNA-LNP encoded membrane-bound immunogen displaying three conserved HIV-1 Envelope (Env) epitopes to knock-in mouse models with B cell receptors (BCRs) of defined affinities, we investigated how epitope-specific competition shapes germinal center (GC) responses. Co-activation of B cells targeting different epitopes did not alter GC kinetics observed in individual activations, but a striking inverse correlation was observed between BCR affinity and GC residence time in either scenario: high-affinity B cells exhibited shorter persistence in GCs, while those with lower affinity to the antigen were maintained. Furthermore, B cells were able to engage in GC reactions at equivalent rates in the presence or absence of clonal lineages binding the same epitope with similar affinities, while higher-affinity clones suppressed lower-affinity counterparts targeting the same epitope. Spatial transcriptomics revealed plasma-like cells within and adjacent to the GC which, together with the detection of early IgG in draining lymph nodes, suggests that local antibody production from these cells may contribute to feedback-driven kinetics. These findings indicate that a self-modulated local antibody feedback loop may act as a “brake” on epitope-specific recognition—dampening further affinity enhancement for high-affinity B cells and facilitating epitope spreading by redirecting the response toward alternative epitopes.

## Introduction

The fundamental process underpinning the efficacy of the humoral response to infection or vaccination is affinity maturation^1,2^. Exposure to antigen activates a polyclonal B cell response derived from those cells with B cell receptors (BCRs) meeting a minimum affinity threshold for the antigen^3^. Those with very high initial affinities will differentiate into extrafollicular plasma cells^4^, while others will form a specialized microanatomical structure, the germinal center (GC). GCs generate and diversify B cells: therein, somatic hypermutation (SHM) introduces mutations into the variable region of BCRs, producing further variation in affinity for the antigen^5^.

This expanded diversity is then shaped by the pressure of competition in GC selection. Expansion in the GC is predicated upon the receipt of survival and proliferation signals from T follicular helper (Tfh) cells^6,7^. B cells engage Tfh by presenting internalized antigen complexed with MHC II^8,9^, and the efficiency of this presentation is affinity dependent^3,10^. Though low affinity clones can enter and participate in GCs, they may be excluded entirely, or simply disfavored for persistence, in the presence of higher affinity competitors^11–13^. Lower affinity B cells surviving in the GC may be selected for differentiation into memory B cells (MBC)^14,15^, but there is some dispute as to whether fate differentiation into MBC and long-lived plasma cell (LLPC) is determined by BCR affinity^16^, and there is substantial variation in the affinity of antibodies expressed by plasma cells emerging from the GC^17^. Whatever the driver of terminal differentiation, slower antigen-BCR dissociation reaches a point of diminishing returns in terms of receptor activation enhancement, suggesting that, even if high affinity responses appear quite early in the response, maturation in the GC ultimately reaches a ceiling after which affinity improvements are no longer visible to selection^3,18,19^. The antibodies of varying affinities produced by terminally differentiated B cells arising throughout this response may also play a role in shaping the interactions of B cells in GCs.

Feedback from existing antibodies has long been known to influence the course of the downstream antibody response^20–23^, but the mechanisms by which it influences the diversity of B cells entering and exiting GCs over the course of the response remains underexplored. Pre-existing antibody allows for immune complex formation^24^, and the BCR affinity activation floor lowers for soluble antigens complexed with antibody^3^, potentially suggesting a relaxation in selection. Conversely, circulating antibody may be a competitive force, intensifying selective pressure in GCs by limiting access to antigen^25^ or blocking cognate B cell entry entirely^26^. Prior work from our own group suggests a circulating Ab affinity tipping point, with lower affinity Abs enhancing GC responses and higher affinity Abs blocking^27^. Furthermore, antibodies influence B cell responses depending on their dose, affinity, and epitope specificity^28^. However, when high affinity mAbs were present prior to SARS-CoV-2 immunization, a preponderance of low affinity clones were observed in post-vaccination GCs^29^; notably, the SARS-CoV-2 booster response was also characterized by a shift to subdominant epitopes. Breadth, and not potency alone, is an important determinant of protection, and a study of malaria vaccine boosting also found that, while Abs from prior vaccination limited overall recall responses, those later responses were also characterized by a shift towards subdominant epitopes^30^. These results highlight the need for further research to delineate the conditions under which antibody feedback constrains or promotes B cell diversity within GCs.

This exploration is particularly important for the development of vaccines requiring a longer immune response, multiple boosts, or multiple epitopes. Those features are characteristic of HIV-1 retrovaccinology based on broadly neutralizing antibodies (bnAbs) to conserved epitopes on the HIV Envelope (Env) isolated from natural infections. In “germline-targeting” (GT) strategies, priming immunogens are designed for high affinity to the unmutated B cells inferred to give rise to bnAbs; these precursors are often relatively rare and lack affinity to the mature HIV trimer^31–42^. After a successful prime, several boosts with sequentially more native-Env-like immunogens may be required to guide the precursors towards functional anti-HIV bnAbs^32,39,41,43^. An effective vaccine will likely need to elicit bnAbs to two or more distinct epitopes on Env to prevent viral escape^44–50^. An individual undergoing these proposed vaccination schedules would thus have a variety of B cell lineages at varying affinities for conserved epitopes on Env and fluctuating antibody titers, converging on the same native trimer. Further complicating this is our novel capacity to introduce membrane-anchored antigen presentations via mRNA-LNPs; increasing avidity by presenting antigen on a surface may lower the threshold for initial activation^51^, or otherwise unpredictably alter the competitive environment.

Navigating cell competition and antibody feedback in the GC environment is therefore crucial to the efficacy of vaccines and, conversely, preclinical work on vaccination provides a unique arena for investigating basic immune biology. To dissect these mechanisms, we established a series of late-stage HIV preclinical mouse models carrying human BCRs of known affinities to one of three epitopes on the HIV-1 Env: the CD4 binding site (CD4bs), V2-Apex, and V3-glycan. These BCRs are “intermediate”—falling between the germline and mature bnAb in terms of affinity and sequence evolution. By vaccinating variable-affinity preclinical B cell models with an mRNA-LNP encoding a single, membrane-anchored native trimer immunogen which presents all three epitopes, we found that, while lower affinity cells reacted robustly to the trimer in isolation, competition from higher affinity lineages could completely block the response of lower affinity lineages to the same epitope. Conversely, lower affinity B cell lineages specific to the same epitope, and lineages with different epitope specificities, did not affect each other’s GC kinetics. The introduction of high affinity antibodies alone was sufficient to reproduce this phenotype, suggesting that antibody feedback underpins this clonal competition; furthermore, an investigation of the local, draining lymph node (dLN) revealed local plasma cells within GCs as a likely source of blocking antibody. These unique “real world” preclinical models not only provide new insights into the mechanisms governing B cells within germinal centers but also offer guiding principles for the rational design of vaccines to elicit potent and broad antibody responses.

## Results

### High-affinity BCR clones are efficiently recruited to germinal centers but are not preferentially maintained over lower-affinity counterparts

To investigate how inherent affinity affects GC kinetics following immunization with HIV-1 Env glycoproteins, we developed mouse models with BCRs derived from the phylogenies of human bnAbs with distinct epitope specificities (**Fig. S1A–C**). We selected clinically-relevant bnAbs targeting three non-overlapping neutralizing epitopes on the highly glycosylated HIV-1 Env: N6-I3 (CD4bs)^52,53^, PCT64-18D (V2-apex)^54^, and minBG18.6 (V3-glycan) (**Fig. S1D–F**)^40^. Using our one-step CRISPR/Cas9-induced homologous directed recombination method^55,56^, we generated knockin (KI) mouse models with B cells expressing the heavy chains (HCs) and/or light chains (LCs) of these three Abs, enabling direct comparison of their GC responses. Lines were bred to homozygosity before use; follicular cell development was normal, and though lines showed some variation in peritoneal B cells all expected populations were present (**Fig. S1G&H**). B cells isolated from these lines survived, differentiated and proliferated at normal rates in vitro (**Fig. S1I**). Peripheral blood mononuclear cells (PBMCs) in the resulting Η^N6-I3/N6-I3^κ^N6-I3/N6-I3^ (referred to as CD4bs-Int^1^ below), Η^PCT64–18D/PCT64–18D^κ^PCT64–18D/PCT64–18D^ (V2-Int^1^), and Η^minBG18.6/minBG18.6^κ^WT/WT^ (V3-Int^1^) KI mouse lines were examined by 10x single-cell RNA sequencing (scRNAseq), revealing that KI HC and LC sequences were co-expressed in 93.1% and 99.7% of cells from CD4bs-Int^1^ and V2-Int^1^ KI mice, respectively, confirming intended KI BCR assembly. For the V3-Int^1^ line, HCs paired with diverse native murine LCs, most frequently IGLV1, IGKV4-68, IGKV4-61, IGKV12-98 and IGKV4-91 (**Fig. 1A**). B cells from these 3 lines were then tested for binding capacity to the HIV Env trimer. We required an immunogen capable of activating B cells targeting diverse epitopes across a broad affinity range to evaluate how inherent BCR affinity impacts GC kinetics. BG505 MD39.3 gp151 (BG505 hereafter), which has performed well in NHPs and a human clinical trial^57,58^, met that criterion. PBMCs from all three lines showed competent specific binding to BG505, and lack of binding to corresponding epitope knockout (KO) probes (**Fig. 1B and Fig.1C**).

**Figure 1.**
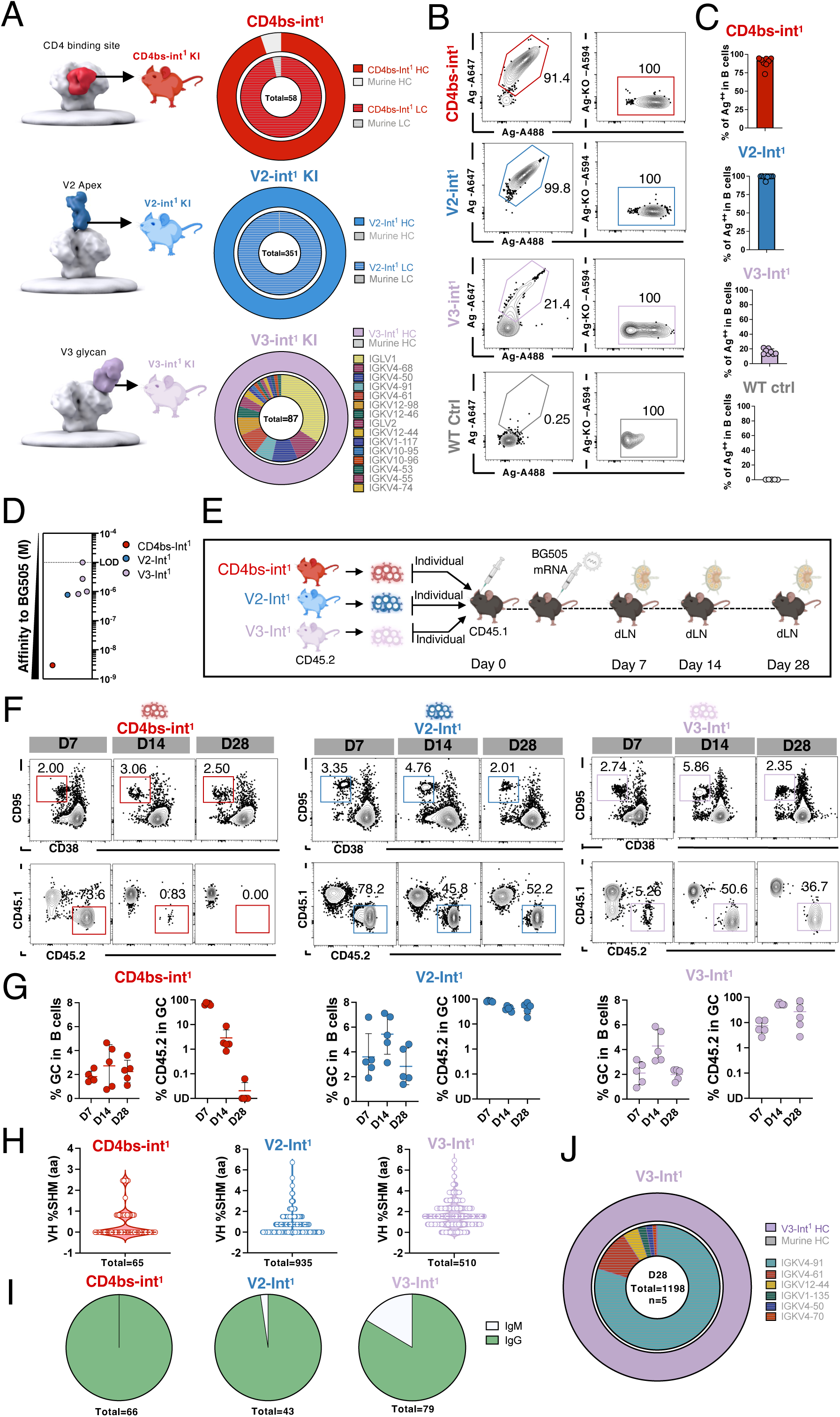
An mRNA-encoded membrane-anchored BG505 native trimer recruits intermediate KI B cells targeting different epitopes on Env. (A) (Top) Nested pie chart of CD4bs-Int^1^ heavy chain (HC) (red), murine HC (gray), human CD4bs-Int^1^ light chain (LC) (shaded red) and murine LC (shaded gray) sequences amplified from single-cell sorted epitope-specific (BG505^+^KO^−^) naive B cells in CD4bs-Int^1^ KI mice. (Middle) As top for V2-Int^1^ HC (blue), murine HC (gray), human V2-Int^1^ LC (shaded blue) and murine LC (shaded gray). (Bottom) As prior for human V3-Int^1^ HC (purple), murine HC (gray) and murine LC sequences (multi-colored). Outer rings represent HCs and inner rings represent LCs. Data are shown from one representative of two exq9ew-ruiopperiments (n=1–2 mice). Total denotes single cells amplified. For V3-Int^1^, the top 15 of 20 LCs amplified are listed. (B) Representative FACS plots of BG505-double-positive and BG505-epitope-specific-KO-negative peripheral B cells in (descending): naïve CD4bs-Int^1^, V2-Int^1^, V3-Int^1^ KI mice, or WT controls. Events were pre-gated on lymphocytes/singlets/CD4^-^CD8^-^F4/80^-^Gr1^-^/B220^+^ B cells. (C) Quantification of (B). BG505-specific binders in blood peripheral B cells in KI mice. n=8–10 for KI mice, n=5 for WT, pooled from 2–4 individual experiments. Bars are mean + SD. (D) Affinity of CD4bs-Int^1^ or V2-Int^1^ or V3-Int^1^ Ab against BG505 trimer measured by SPR dissociation constant. For V3-Int^1^, Abs were expressed with human HC and representative murine LCs, detailed sequences can be found in the key resource table. Each dot represents the mean of 4 technical replicates. Dotted line marks the limit of detection (LOD). (E) Schematic of GC recruitment evaluation experiments for CD4bs-Int^1^, V2-Int^1^, or V3-Int^1^ KI B cells. Immunization output panels below display representative data from one of at least two experiments with 3–5 mice per condition. (F) Representative FACS plots of B cells obtained from dLNs after immunization with mRNA-LNP encoding BG505 Env glycoprotein trimer. Days post-immunization at top. Events pre-gated on lymphocytes/singlets/live/CD4^-^CD8^-^F4/80^-^Gr1^-^/B220^+^ B cells and represent GC in B cells, or CD45.2 cells in GC. (G) Time course plots of GC cells as a percentage of total B cells (left) and CD45.2^+^ KI cells as a percentage of GC B cells (right). Values of zero were plotted as UD (undetected) on the log₁₀ scale. Each dot represents one mouse. Bars are mean ± SD. (H) Dotted violin plots of HC amino acid mutations across all sites at day 21. Each dot represents an HC sequence from one B cell. (I) Pie charts of class-switch profiles at day 21. Total = number of sequences amplified. (J) Nested pie chart showing V3-Int^1^ HC and LC usage from single-cell sorted epitope-specific (BG505^+^KO^−^) B cells at day 28; no murine HC (grey) detected. Outer layer, human V3-Int^1^ IGHV; inner layer, murine IGKV. Nested pie chart shows average of all mice in a group (Total=number of sequences amplified, n=number of mice).

All three BCR variants showed detectable but varied affinities to BG505: the dissociation constant [*K*_D_s] of CD4bs-Int^1^ was 3 nM, V2-Int^1^ 770 nM, and V3-Int^1^, where the minBG18.6 heavy chain paired with representative mouse light chains, ranged from 820 nM to the limit of detection (LOD) at 10 µM (**Fig. 1D**). CD4bs-Int^1^, V2-Int^1^, or V3-Int^1^ CD45.2^+/+^ KI cells were adoptively transferred into WT CD45.1^+/+^ hosts to reach a combined frequency of 20 in 10^6^, and recipient mice were then immunized with an mRNA-LNP-encoded membrane-anchored BG505 (**Fig. 1E**). The GC response and the proportion of CD45.2^+^ B cells in GCs was evaluated in dLNs at days 7, 14, and 28. Across all transfer recipients, GCs constituted on average ∼2–5% of total B cells throughout (**Fig. 1F and Fig. 1G**). CD45.2^+^ KI cells of each line were effectively activated and present in GCs at day 7, though the average fraction of CD45.2^+^ cells in GCs ranged from 5% for V3-Int^1^ to 67% for CD4bs-Int^1^ and 78% for V2-Int^1^. At day 14 and day 28, V2-Int^1^ and V3-Int^1^ CD45.2 B cells retained a share of the GC (43% and 47% for V2-Int^1^, 53% and 27% for V3-Int^1^); V3-Int^1^ responses displayed higher variability at day 28, consistent with the polyclonality of naïve V3-Int^1^ BCRs. In contrast to either V2-Int^1^ or V3-Int^1^, CD4bs-Int^1^ dropped to 2.88% by day 14 and 0.16% at day 28—notably, CD4bs-Int^1^ cells have the highest affinity for their epitope (*K*_D_, 3nM) (**Fig. 1F and Fig. 1G**). Sequencing by single-cell BCRseq of sorted antigen-specific CD45.2^+^ B cells at day 21 revealed that immunization stimulated class-switching and SHM in all three cell lines **(Fig. 1H**, **Fig. 1I, Fig. S1J)**. Immunization selected for particular V3-Int^1^ BCRs, with a notable shift towards IGKV4-91 (79.88%) and IGKV4-61 (10.93%) (**Fig. 1A, and Fig. 1J**). Thus, independent of initial affinity, all three cell lines could be stimulated by an mRNA-LNP-delivered membrane-anchored immunogen, enter the early GC, and undergo class switching and SHM, but each line displayed distinct GC kinetics, with shorter GC half-lives observed for the line with highest initial affinity to BG505.

### High-affinity clones do not influence GC dynamics of low-affinity B cells targeting other epitopes

Considering the variation in affinity across these three knockin lines (**Fig. 1D)**, we next sought to determine whether they would compete despite targeting different epitopes on the same trimer. Therefore, CD4bs-Int^1^, V2-Int^1^, and V3-Int^1^ CD45.2^+/+^ KI B cells were adoptively transferred into WT CD45.1^+/+^ mice individually to reach a frequency of 20 cells per 10^6^ total B cells, or in equal-proportion combinations to reach 20 (1x Mix) or 60 (3x Mix) cells per 10^6^ total B cell. 1x Mix matched the total KI fraction to that in individual transfers while in the 3x Mix condition individual KI lines were at frequencies obtained after individual transfers but together made up a larger fraction of the compartment. Recipient mice were immunized intramuscularly with 2 µg of BG505 mRNA-LNP one day later. Local, draining LNs were analyzed by flow cytometry at days 7 and 21 post-immunization (**Fig. 2A**). GC sizes were similar across groups at both timepoints; V2-Int^1^ alone produced slightly larger GCs at day 7 than 3x Mix, but this difference did not persist to 21 (**Fig. 2B**–**2C**). Mice transferred with V3-Int^1^ alone averaged fewer CD45.2^+^ B cells in GCs at day 7 relative to recipients of other individual transfers or either Mix, but no significant differences were observed at day 21 (**Fig. 2D**). To determine the composition of the CD45.2 B cells in the mixed transfer recipients, all CD45.2 KI cells in the GCs were sorted for 10x scRNA-seq. CD4bs-Int^1^, V2-Int^1^, and V3-Int^1^ cells were recovered from both the 1x and 3xMix recipients at variable frequencies (**Fig. 2E**). The ratio for each KI cell identified by scRNAseq was multiplexed back to the total CD45.2 KI cell percentage obtained by flow cytometry for each individual mouse to determine the estimated specific single KI cell rates in dLN B cells. The percentage of CD4bs-Int^1^, V2-Int^1^, and V3-Int^1^ B cells among total B cells at day 21 was indistinguishable between single transfers and both combination transfers (1x and 3x dose) (**Fig. 2F, Fig. S2A**). Slightly higher mutation rates were observed in all three B cell lines isolated from 1x Mix recipients compared to individual transfer recipients; in 3x Mix recipients, SHM was only increased for V3-Int^1^ HCs (**Fig. S2B**). No differences were observed in class-switching profiles, or the identities of the three most common LC found with V3-Int^1^, though frequencies of these LCs varied by treatment (**Fig. S2C–D**). The GC kinetics of each epitope-specific cell line was thus unaffected by the presence of lines targeting non-overlapping epitopes on the same immunogen.

**Figure 2.**
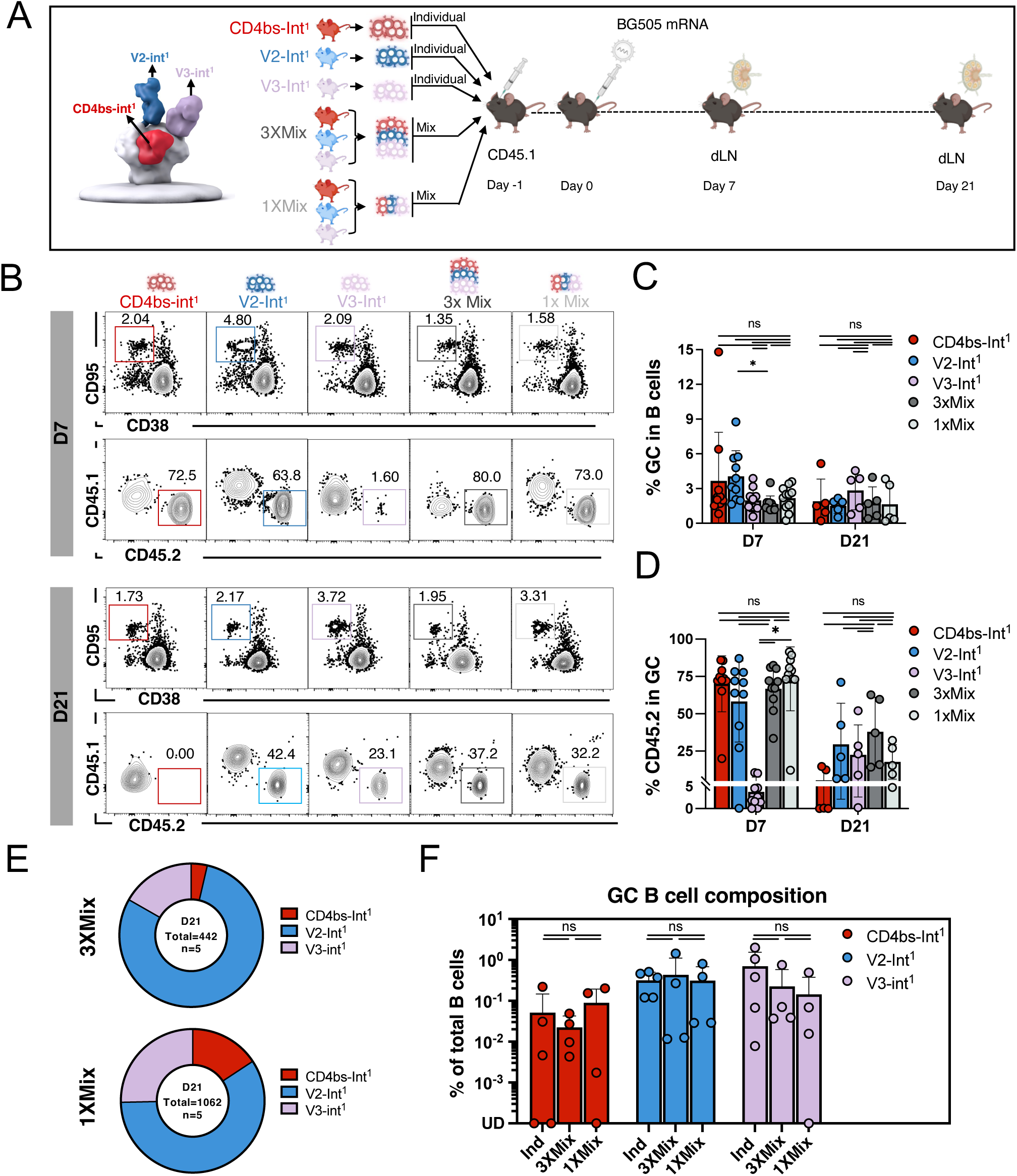
A single trimer can simultaneously activate intermediate B cells targeting V2-apex, V3-glycan, and the CD4bs in the same host. (A) Schematic of individual- or co-adoptive-transfer and immunization experiments for CD4bs-Int^1^, V2-Int^1^, and V3-Int^1^ KI B cells. (B) Representative FACS plots of B cells obtained from dLNs after individual- or co-adoptive-transfer. Events pre-gated on lymphocytes/singlets/live/CD4^-^CD8^-^F4/80^-^Gr1^-^/B220^+^ B cells and represent (upper) GC and (lower) CD45.2 cells in GC. (C–D) Quantification of (C) GC cells as the percentage of total B cells and (D) CD45.2^+^ KI cells as the percentage of GC B cells at days 7 and 21. Adjusted p-values (q-values) were calculated by Kruskal-Wallis test followed by pairwise comparisons with Benjamini–Krieger–Yekutieli (BKY) correction for multiple comparisons. (E) CD4bs-Int^1^ (red), V2-Int^1^ (blue), and V3-Int^1^ (purple) B cell lineage frequency analysis from 10x scRNAseq of GC CD45.2 B cells sorted at day 21 from the 3x Mix (top) and 1x Mix (bottom) groups. Pie charts are averages of all mice in a group (Total=number of sequences amplified, n=number of mice). (F) KI BCR composition of GC CD45.2 B cell sequences as percentages of total B cells at day 21. Percentage was calculated from FACS data for individual groups, and from 10x scRNAseq plus FACS data for Mix groups. Values of zero on the log₁₀ scale were plotted as UD. Adjusted p-values (q-values) were calculated by 2-way ANOVA test followed by pairwise comparisons with BKY correction for multiple comparisons. Figures represent data from one of at least two experiments with 3–5 mice per condition. Bars are mean + SD. Each dot represents one mouse. Adjusted p-values (q-values) were indicated as follows: *p < 0.05, ns = not significant.

### B cell lines with low, overlapping affinities to the same epitope can be co-stimulated

Although we did not observe competitive inhibition among B cell lines targeting different epitopes on the same mRNA-LNP-encoded immunogen, to explore the possible crosstalk between lineages targeting the same epitope, we generated an additional HC KI mouse line using minBG18.11 (V3-Int^2^), which targets the same V3-glycan epitope as minBG18.6 (V3-Int^1^) but represent a more affinity-matured variant with higher SHM from germline^40^. Homozygous V3-Int^2^ mice (Η^minBG18.11/minBG18.11^κ^WT/WT^) displayed normally developing B cells which can survive, proliferate, and differentiate in vitro (**Fig. S3A–C**). BG505-specific PBMC frequency was 27.9%, similar to that observed in V3-Int^1^ (**Fig. 3A, Fig. S3D, Fig. 1B–C**). As with V3-Int^1^, the KI V3-Int^2^ HC sequences also paired with a variety of endogenous murine LCs, most commonly IGKV12-46, IGKV12-41, IGKV10-94, IGKV4-50, and IGKV12-98 (**Fig. 3B**). V3-Int^1^ and V3-Int^2^ displayed similar GC kinetics post-immunization by BG505 mRNA-LNP, with V3-Int^2^ showing a slightly decreased GC size from 3.2% to 1.5% between D14 and D28; CD45.2^+^ cells in GC were 35% at day 14 and 45% at day 28 (**Fig. 3C–D, Fig. S3E–F**). While V3-epitope recognition is strongly HC dependent^40^, the recombination with murine LCs adds further diversity and the affinity of monoclonal antibodies (mAbs) representative of V3-Int^2^ BCRs to BG505 ranged from 1 µM to 10 µM, comparable to V3-Int^1^ representative mAbs (**Fig. 3E**).

**Figure 3.**
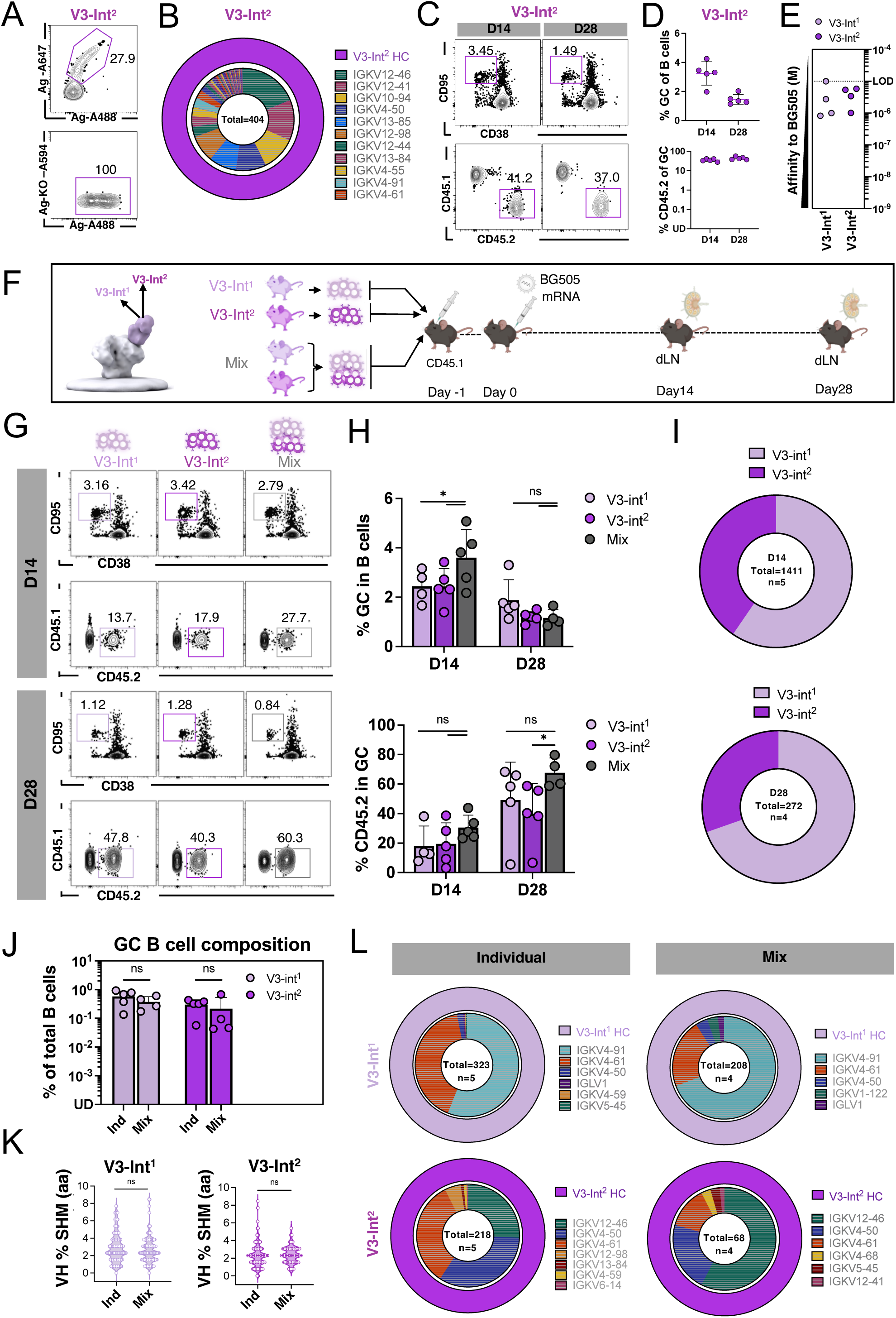
Two KI lineages targeting V3-glycan can be simultaneously activated without competitive interference. (A) Representative FACS plots of BG505-double-positive and BG505-epitope specific KO-negative peripheral B cells in naïve V3-Int^2^ KI mice. Events were pre-gated on lymphocytes/singlets/CD4^-^CD8^-^F4/80^-^Gr1^-^/B220^+^ B cells. (B) Nested pie chart of human V3-Int^2^ HC (purple), murine HC (gray) and murine LC sequences (multi-colored) amplified from single-cell sorted epitope-specific (BG505^+^KO^−^) naive B cells in V3-Int^2^ KI mice. Legend shows names of most frequent 11 of 43 LCs amplified. Outer rings represent HCs and inner rings represent LCs. Total = single cells amplified. (C) Representative FACS plots of B cells obtained from dLNs in V3-Int^2^ adoptively transferred mice at days 14 and 28 points after immunization with mRNA-LNP-encoded BG505. Events pre-gated on lymphocytes/singlets/live/CD4^-^CD8^-^F4/80^-^Gr1^-^/B220^+^ B cells and represent GC, CD45.2 cells in GC. (D) Quantification of (C). Each dot represents one mouse. Bars are mean ± SD. (E) Affinity measurement of V3-Int^1^ and V3-Int^2^ Ab against BG505 trimer measured by SPR dissociation constant. V3-Int^1^ and V3-Int^2^ Abs were expressed with human HC and representative murine LCs, detailed sequence can be found in the key resource table. Each dot represents the mean of four technical replicates. Dotted line marks LOD. Data for V3-Int^1^ reproduced from Fig. 1D. (F) Schematic of individual or co-adoptive-transfer and immunization experiments for V3-Int^1^ and V3-Int^2^ KI B cells. (G) Representative FACS plots of B cells obtained from dLNs of mice V3-Int^1^ and V3-Int^2^ B cells at days 14 and 28 of BG505. Events were pre-gated on lymphocytes/singlets/live/CD4-CD8-F4/80-Gr1^-^/B220^+^ B cells and represent GC and CD45.2 cells in GC. (H) (Upper) GC cells as a percentage of total B cells and (lower) CD45.2^+^ KI cells as a percentage of total GC B cells at days 14 and 28. Each dot represents one mouse. Adjusted p-values (q-values) calculated by 2-way ANOVA test with pairwise post-hoc comparisons adjusted using BYK correction. (I) The V3-Int^1^ (light purple) and V3-Int^2^ (dark purple) B cell lineage frequency analysis from 10x scRNAseq of GC CD45.2 B cells sorted from the Mix group. Pie charts are averages of all mice in a group (Total=number of sequences amplified, n=number of mice). (J) KI BCR composition of GC CD45.2 B cell sequences as a percentage of total B cells. Percentage was calculated from FACS data for individual groups, and from 10x scRNAseq plus FACS data for Mix groups. Values of zero were plotted as UD on the log₁₀ scale. Adjusted p-values (q values) calculated by 2-way ANOVA Kruskal-Wallis test with pairwise post-hoc comparisons adjusted using BYK correction. (K) Dotted violin plot of HC amino acid mutations across all sites at day 28. Each dot represents an HC sequence from one B cell. Statistical analysis made using Mann-Whitney’s test. (L) (Upper) Nested pie chart showing V3-Int^1^ HC and LC usage from single-cell sorted CD45.2 B cells at day 28 from individual transfer and mix recipients. Outer ring, human V3-Int^1^ IGHV; inner ring, murine IGKV. (Lower) Nested pie chart showing V3-Int^2^ HC and LC usage as upper. Nested pie charts are averages of all mice in a group (Total=number of sequences amplified, n=number of mice). Panels in C, D, G, H display representative data from one of at least two experiments with 3–5 mice per condition. Bars are mean ± SD in (H) and (J). Adjusted p-values (q-values) are indicated as follows: *p < 0.05, **p < 0.01, ***p < 0.001, ****p < 0.0001; ns = not significant.

To establish a repertoire of B cells potentially competitive for the same epitope, V3-Int^2^ and V3-Int^1^ CD45.2^+/+^ B cells were adoptively transferred into CD45.1^+/+^ host mice either individually, to reach a combined frequency of 20 cells per 10^6^ total B cells, or in equal-proportion combinations (mix) to reach a combined frequency of 40 cells per 10^6^ total B cells. Recipients were *i.m.* immunized one day later (day 0) with BG505 mRNA-LNP, and local dLNs were analyzed at days 14 and 28 (**Fig. 3F**). Mix-transfer recipients had larger GCs than either single-transfer group at day 14, but no significant differences were observed at day 28 (**Fig. 3G–H**). The frequency of antigen-specific CD45.2^+^ B cells in GCs increased over time. At day 28, a slightly higher frequency of CD45.2^+^ B cells was observed in GCs in the mix-recipients compared to V3-Int^2^-recipients, but frequencies were otherwise similar across groups at both time points (**Fig. 3G–H**). CD45.2^+^ KI cells from mix-transfer recipients were also sorted for 10x scRNAseq; both V3-Int^1^ and V3-Int^2^ B cells were recovered at days 14 and 28 (**Fig. 3I, Fig. S3G**). Upon multiplexing the ratios back to the GC CD45.2^+^ KI cell percentages per mouse, no significant difference in relative B cell frequencies was observed between the single-transfer and the mix-transfer groups (**Fig. 3J, Fig.S3H**). While V3-Int^1^ HC isolated from mix recipients had higher SHM at D14 than those from individual transfer groups, these differences equilibrated by day 28 and no differences were apparent in V3-Int^2^ HC SHM across treatments (**Fig. 3K, Fig. S3I**). V3-Int^1^ HC maintained their preference for IGKV4-91, IGKV4-61, and IGKV4-50 LCs whether developing alone or accompanying V3-Int^2^; similarly, V3-Int^2^ HC prefers IGKV12-46, IGKV4-50, and IGKV4-61 in both single and mixed transfer scenarios, though the frequencies of these most common pairings varied (**Fig. 3L**). We observed no differences in class-switching profiles for either V3-Int^1^ or V3-Int^2^ B cells between individual and mix-transfer recipients (**Fig. S3J**). Thus, despite engaging precisely the same epitope on the mRNA-LNP-delivered immunogen with overlapping affinity ranges, V3-Int^1^- and V3-Int^2^-derived B cells do not alter one another’s GC participation or SHM accumulation.

### High-affinity B cells filter out lower-affinity cells targeting the same epitope

As V3-Int^1^ and V3-Int^2^ have variable, and overlapping, affinities to BG505, to further clarify the effects of affinity on cells competing for the same epitope, we generated an additional HC+LC line targeting the CD4bs. We used the sequences of the Ab min12A21^53^ to generate Η^min12A21/min12A21^κ^min12A21/min12A21^ (referred to as CD4bs-Int^2^ below), which has lower levels of SHM than CD4bs-Int^1^. The affinity of CD4bs-Int^2^ to the BG505 trimer (46 nM) is lower than that of CD4bs-Int^1^ (3 nM) (**Fig. 4A**), though both are quite high compared to V3-Int^1^ and V3-Int^2^, which ranged between 1–10 µM. (**Fig. 3E**). An average of 31.2% of CD4bs-Int^2^ KI PBMCs bound the BG505 probe (**Fig. 4B, Fig. S4A**), and 10x scRNAseq demonstrated that the BCRs of binders almost exclusively expressed the full length CD4bs-Int^2^ HC and LC sequences (95% HC and 97% LC), similar to CD4bs-Int^1^ (**Fig. 4C**, **Fig.1A**). CD4bs-Int^2^ development to follicular B cells, as well as survival, proliferation and differentiation in vitro, were comparable to CD4bs-Int^1^ (**Fig. S4B–D**). Following stimulation with BG505 mRNA-LNP, CD4bs-Int^2^ B cells could be recruited to GCs at day 7 and maintained, averaging 43% at day 7 and 29% at day 28 (**Fig. 4D**). This contrasts sharply with CD4bs-Int^1^ B cells, which were nearly undetectable in GCs at day 21 (**Fig. 1G**).

**Figure 4.**
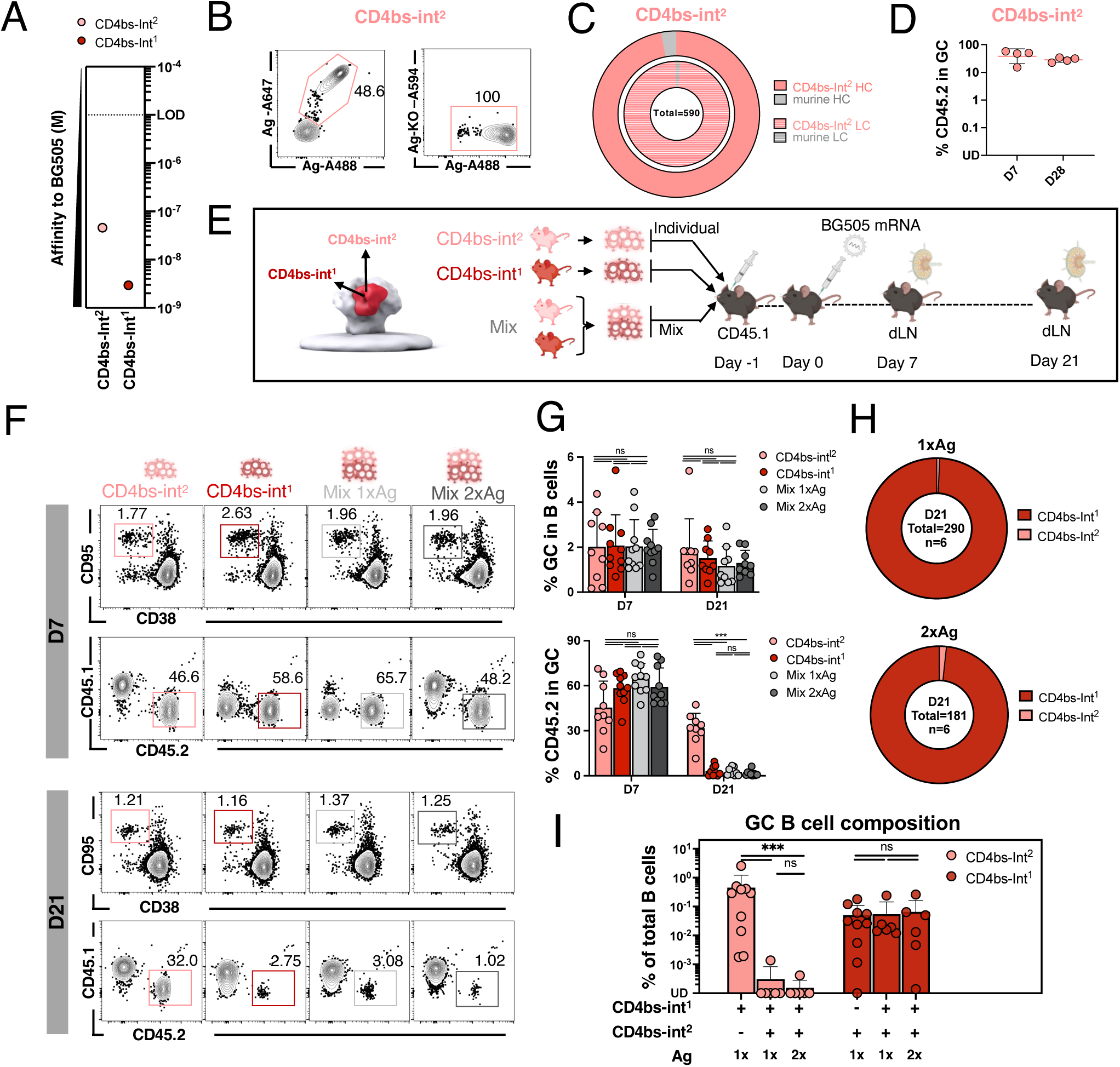
Affinity-dependent inhibition of GC recruitment occurs between KI B cell lines competing for the CD4bs. (A) Affinity of CD4bs-Int^1^ and CD4bs-Int^2^ human Ab against BG505 trimer measured by SPR dissociation constant. Each dot represents the mean of four technical replicates. Dotted line represents LOD. (B) Representative FACS plots of BG505-double-positive and BG505-epitope specific KO-negative peripheral B cells in naïve CD4bs-Int^2^ KI mice. Events were pre-gated on lymphocytes/singlets/CD4^-^CD8^-^F4/80^-^Gr1^-^/B220^+^ B cells. Data is representative of a total of ten mice from five individual experiments. (C) Nested pie chart of human CD4bs-Int^2^ HC (shaded pink), murine HC (shaded gray), CD4bs-Int^2^ LC (pink), and murine LC sequences (gray) amplified from single-cell sorted epitope-specific (BG505+KO−) naive B cells in CD4bs-Int^2^ KI mice. Total = single cells amplified. Data were produced from one representative of three mice. (D) CD45.2^+^ KI cells as a percentage of total GC B cells in dLNs of CD4bs-Int^2^ adoptive transfer recipients at 7 and 21 days post-immunization by BG505. Values of zero were plotted as UD on the log₁₀ scale. Each dot represents one mouse. Bars are mean ± SD. (E) Schematic of individual or co-adoptive-transfer and immunization experiments for CD4bs-Int^1^ and CD4bs-Int^2^ KI B cells. Recipient mice were adoptively transferred with either CD4bs-Int^1^ or CD4bs-Int^2^ KI B cells to reach a frequency of 20 in 10^6^ B cells before immunization with 2 µg of BG505 mRNA. Mice were sacrificed at day 7 and day 21. (F) Representative FACS plots of B cells obtained from dLNs at days 7 and 21 post-immunization by BG505 in individual or mix transfer recipients. Events were pre-gated on lymphocytes/singlets/live/CD4^-^CD8^-^F4/80^-^Gr1^-^/B220^+^ B cells and represent GC, CD45.2 cells in GC. (G) (upper) GC cells as a percentage of total B cells and (lower) CD45.2^+^ KI cells as a percentage of total GC B cells at days 7 and 21. (H) The CD4bs-Int^2^ (pink) and CD4bs-Int^1^ (red) B cell lineage frequency analysis from 10x scRNAseq of GC CD45.2 B cells sorted from the Mix groups at day 21. Pie charts are averages of all mice in a group (Total=number of sequences amplified, n=number of mice). (I) BCR composition of GC CD45.2 B cell sequences as a percentage of total B cells at day 21. Percentage was calculated from FACS data for individual groups, and from 10x scRNAseq plus FACS data for Mix groups. Values of zero were plotted as UD on the log₁₀ scale. Panel G–I present data pooled from 2 experiments of 3–5 mice per condition. Bars are mean ± SD in (G) and (I). Each dot represents one mouse. Adjusted p-values (q-values) calculated by nonparametric Kruskal-Wallis test with pairwise post-hoc comparisons adjusted using BYK correction are indicated as follows: ***p < 0.001; ns = not significant.

After demonstrating lack of inhibitory competition between V3 cells with low, overlapping-affinity ranges, the CD4bs-Int^2^ and CD4bs-Int^1^ KIs allow for the exploration of same-epitope B cells with high affinities differing by approximately 15-fold. CD4bs-Int^1^ and CD4bs-Int^2^ CD45.2^+^ B cells were adoptively transferred into WT mice either individually (to achieve a frequency of 20 per 10^6^ total B cells) or in equal combination (to achieve a combined frequency of 40 per 10^6^ total B cells), and recipients immunized with BG505 mRNA-LNP (**Fig. 4E**). All groups displayed similar GC sizes (**Fig. 4F&G, upper**). At day 7, CD45.2^+^ B cell frequencies in GCs were comparable across single and mixed transfer recipients. By day 21, however, CD4bs-Int^2^ B cells maintained high GC presence only when transferred individually, with significantly reduced frequencies observed when co-transferred with high-affinity CD4bs-Int^1^ B cells (**Fig. 4F&G, lower**). In contrast, the frequency of high-affinity CD4bs-Int^1^ B cells in GCs decayed rapidly to nearly undetectable levels at day 21 in both single and mixed transfer recipients (**Fig. 4F&G, lower**). Doubling the antigen dose did not rescue the CD4bs-Int^2^ response (**Fig. 4E–G**).

In mix-transfer recipients, scRNAseq of CD45.2^+^ GC B cells revealed that the higher affinity CD4bs-Int^1^ B cells dominated GCs at both D7 and D21 independent of administered antigen dose, with only ∼5–10% of CD45.2^+^ GC B cells belonging to the CD4bs-Int^2^ lineage (**Fig. 4H, Fig. S4E&F**). As above, we multiplexed the ratio to make the comparison per mouse and found a 1000-fold decrease of CD4bs-Int^2^ cells in GCs after the addition of CD4bs-Int^1^; CD4bs-Int^1^ numbers, in contrast, were unaffected by CD4bs-Int^2^ (**Fig. 4I**), demonstrating inhibition of CD4bs-Int^2^ recruitment to or retention in GCs in the presence of CD4bs-Int^1^. While CD4bs-Int^2^ HC underwent lower rates of mutation in mixture recipients than when transferred alone, the CD4bs-Int^1^ mutation rate was not affected by the presence of the low affinity CD4bs-Int^2^ line (46 nM) (**Fig. S4G**). No differences in class-switch profiles were observed in either line (**Fig. S4H**).

To determine whether the competition for GC residence was mediated by inherent BCR affinity to the administered antigen, we generated a third CD4bs-targeting KI line, Η^N6-I2/N6-I2^κ^N6-I2/N6-I2^, referred to as CD4bs-Int^3^, from another CD4bs targeting Ab: N6-I2^52^, with affinity for the BG505 MD39 antigen (*K*_D_, 5.7 nM) that was only slightly lower than the CD4bs-Int^1^ affinity (*K*_D_, 3 nM) (**Fig. S5A**). Though CD4bs-Int^3^, like CD4bs-Int^1^, was less prevalent than CD4bs-Int^2^ cells when transferred alone (**Fig. S5B–D**), frequency within CD45.2^+^ GC B cells was in order of descending affinity to BG505 in the triple mix (**Fig. S5E**). The addition of CD4bs-Int^3^ to the transfer-mix did not affect the kinetics of CD4bs-Int^1^ (**Fig. S5F**), though greater SHM was observed in CD4bs-Int^1^ (3 nM) after the addition of similarly high affinity CD4bs-Int^3^ (5.7 nM) cells in the triple mixture (**Fig. S5G**). Thus, while no inhibition was observed among B cell lineages with overlapping micromolar affinities targeting the V3-glycan supersite, same-epitope B cells with higher, nanomolar-range affinities targeting the CD4bs suppressed their lower-affinity counterparts.

### Lowering antigen affinity extends GC residence

The absence of CD4bs-Int^2^ in later GCs in hosts also containing higher-affinity CD4bs-Int^1^ cells targeting the same epitope (**Fig. 4F–-I**), as well as the limited time in GCs for CD4bs-Int^1^ after immunization (**Fig. 1F**), led us to examine more directly the effects of BCR affinity to the antigen on GC kinetics using a more traditional protein immunization. We first performed immunization in mice transferred individually with CD4bs-Int^1^ or CD4bs-Int^2^ using BG505 protein adjuvanted with saponin/MPLA nanoparticles (SMNP)^59^, examining intermediate GC developmental phases (**Fig. 5A&**B). Day 8 GCs were similar after either transfer (2–3%); GCs in lower-affinity CD4bs-Int^2^ recipients peaked at day 16 at 6% and returned to 2% at day 21, while GCs in recipients of the higher affinity CD4bs-Int^1^ remained at 2.5% on day 16 before dropping to 0.7% on day 21 (**Fig. 5C&**D). Within the GC, both CD4bs-Int^1^ and CD4bs-Int^2^ comprised approximately 6–7% of GC B cells at day 8, demonstrating successful recruitment. At day 16, however, GC occupancy rates of the lower affinity CD4bs-Int^2^ increased to approximately 12% and then to 16% on day 21; in contrast, by day 16 the higher affinity CD4bs-Int^1^ cells underwent a massive contraction to 0.4%, from which they did not recover on day 21 (**Fig. 5C&E**).

**Figure 5.**
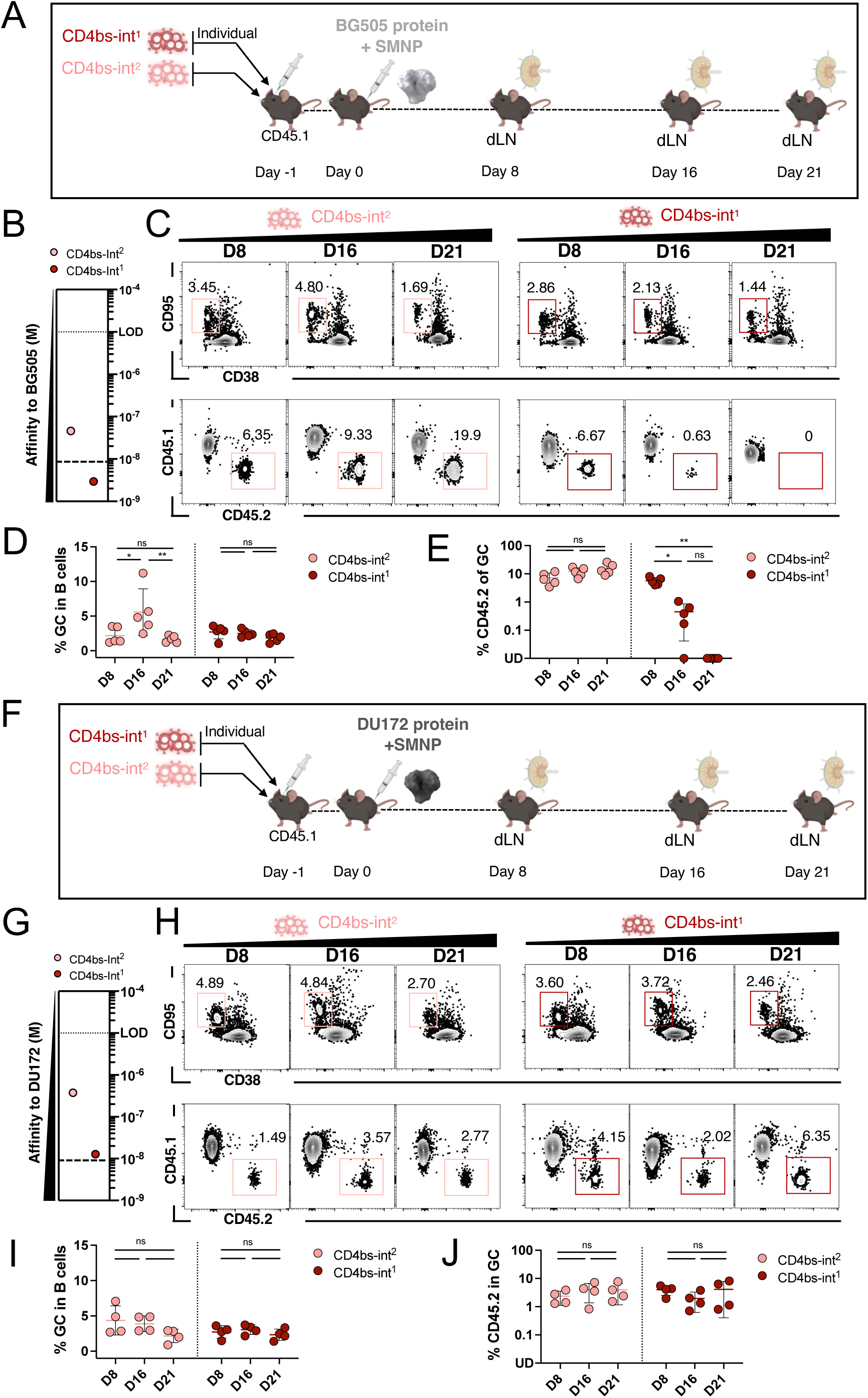
Affinity-related GC kinetics of competing CD4bs-targeting precursor lines. (A) Schematic of BG505 immunization experiments for CD4bs-Int^1^ and CD4bs-Int^2^ B cells. Mice were immunized with BG505 Env glycoprotein trimer protein with SMNP adjuvant. (B) Affinity of CD4bs-Int^1^ and CD4bs-Int^2^ human Ab against BG505 trimer measured by SPR dissociation constant. Each dot represents the mean of 4 technical replicates. Dotted line represents LOD; dashed line proposes a potential affinity selection ceiling in our system in a low nanomolar range. Affinity data reproduced from Fig. 4D for ease of reference. (C) Representative FACS plots of GC B cells obtained from dLNs after BG505 immunization at days 8, 16, and 21. Events were pre-gated on lymphocytes/singlets/live/CD4^-^CD8^-^F4/80^-^Gr1^-^ /B220^+^/GC B cells and represent CD45.2 KI cells in GC. (D) Kinetics of GCs over time after BG505 immunization. (E) Kinetics of CD45.2 cells in GCs over time after BG505 immunization. Values of zero were plotted as UD on the log₁₀ scale. (F) Schematic of DU172 immunization experiments for CD4bs-Int^2^ and CD4bs-Int^1^ KI B cells. Mice were immunized with DU172 Env glycoprotein trimer protein with SMNP adjuvant. (G) Affinity of CD4bs-Int^1^ and CD4bs-Int^2^ human Ab against DU172 trimer measured by SPR dissociation constant. Each dot represents the mean of 4 technical replicates. Dotted line represents LOD, dashed line proposes the potential affinity selection ceiling in a low nanomolar range. (H) Representative FACS plots of GC B cells obtained from dLNs after DU172 immunization at days 8, 16, and 21. Events were pre-gated on lymphocytes/singlets/live/CD4^-^CD8^-^F4/80-Gr1^-^ /B220^+^/GC B cells and represent CD45.2 KI cells in GC. (I) Kinetics of GC B cells after DU172 immunization. (J) Kinetics of CD45.2 KI cells as the percentage of GC B cells after DU172 immunization. Values of zero were plotted as UD on the log₁₀ scale. Panels show data from one representative experiment of at least two performed, with 3–5 mice per condition. Dots represent single mice. Bars are mean ± SD. Adjusted p-values (q-values) calculated by nonparametric Kruskal-Wallis test followed by pairwise post-hoc comparisons adjusted using BKY correction for multiple comparisons. Adjusted p-values (q-values) are indicated as follows: *p < 0.05, **p < 0.01, ns = not significant.

To determine the effect of BCR affinity for different antigens on these kinetics, we then deployed another HIV protein trimer adjuvanted with SMNP, DU172 (**Fig. S6A-D**), which displays lower affinity to both CD4bs-Int^1^ (*K*_D_, 13 nM for DU172 vs 3 nM for BG505) and CD4bs-Int^2^ (*K*_D_, 375 nM for DU172 vs 46 nM for BG505) (**Fig. 5F&G**). Total GC sizes were similar in both lines throughout (∼2–5%) (**Fig. 5H&I**). Strikingly, no significant difference in CD45.2 cell presence in GCs was observed between these lines (**Fig. 5H&J)**, despite the fact that the ratio of CD4bs-Int^1^:CD4bs-Int^2^ BCR affinities for DU172, or the “affinity gap,” is 29.4 folds, greater than their 15.6-fold affinity gap for BG505 (**Fig. 5B&G**). Both CD4bs-Int^1^ and CD4bs-Int^2^ were present in GCs until day 21 (**Fig. 5H&J**). Thus, with BG505 immunization, CD4bs-Int^1^ and CD4bs-Int^2^ were recruited to GCs at day 8 at equivalent rates, but the high-affinity CD4bs-Int^1^ (*K*_D_, 3nM for BG505) B cells diminished by day 16 and were absent at day 21. In contrast, after immunization with the lower affinity antigen DU172, both CD4bs-Int^1^ and CD4bs-Int^2^ remained in GCs over time (out to day 21). Thus, lowering antigen affinity below the ∼3–12 nM ceiling prolongs GC residence, consistent with excessive affinity accelerating clonal contraction.

### Antibody-mediated epitope masking suppresses competing B cells in an affinity-dependent manner

Since B cells targeting the same epitope did not directly compete for antigen binding, while lower-affinity CD4bs-specific B cells seemed to be out-competed in GCs by CD4bs-Int^1^ (**Fig. 4G–I**), we investigated antibody-mediated epitope masking as a potential mechanism to determine whether antibody secretion after early fate determination of high-affinity CD4bs-int^1^ B cells might inhibit B cells specific to the same epitope. First, we established a model with higher-affinity passively transferred Abs and lower-affinity B cells targeting the same epitope (**Fig. 6A**): CD45.2^+/+^ B cells from CD4bs-Int^2^ KI donor mice (46 nM affinity to BG505) were adoptively transferred into CD45.1^+/+^ host mice through the retro orbital sinus (day -1); approximately 8 hours later (timepoint referred to as day -0.5), CD4bs-Int^1^ Abs (3 nM affinity to BG505) were injected into host mice through the tail vein; approximately 16 hours later (day 0), recipient mice were immunized with BG505 mRNA. At day 10 post-immunization, dLNs were sampled for flow cytometry (**Fig. 6A**). Although GCs formed and were a larger fraction of B cells at the higher mAb doses (3 µg and 30 µg), all CD4bs-Int^1^ mAb doses higher than 0.05 µg diminished the percentage of CD4bs-Int^2^ B cells in GCs substantially relative to a control mAb; 0.3 µg decreased CD4bs-Int^2^ B cell participation from 14% to 3%; and at 3 µg and 30 µg, CD4bs-Int^2^ B cells were almost entirely blocked from GCs (1% for 3 µg, 0.3% for 30 µg) (**Fig. 6B–C**). Next, we repeated the passive transfer of CD4bs-Int^1^ Ab followed by immunization in mice instead adoptively transferred with equivalently high affinity CD4bs-Int^1^ B cells (**Fig. 6D**). GC size increases were not significant after mAb delivery, and no significant change in the proportion of CD4bs-Int^1^ B cells in GCs was observed at 0.3 µg compared to the control group; however, near-total blocking was observed when the Ab dose was increased to 3 µg or 30 µg (**Fig. 6E–F**). Thus, low concentrations (0.3 µg) of high affinity CD4bs-Int^1^ mAb were sufficient to inhibit lower-affinity CD4bs-Int^2^ cells, but higher concentrations (3 µg and 30 µg) also inhibited cells of equivalent affinity for the antigen.

**Figure 6.**
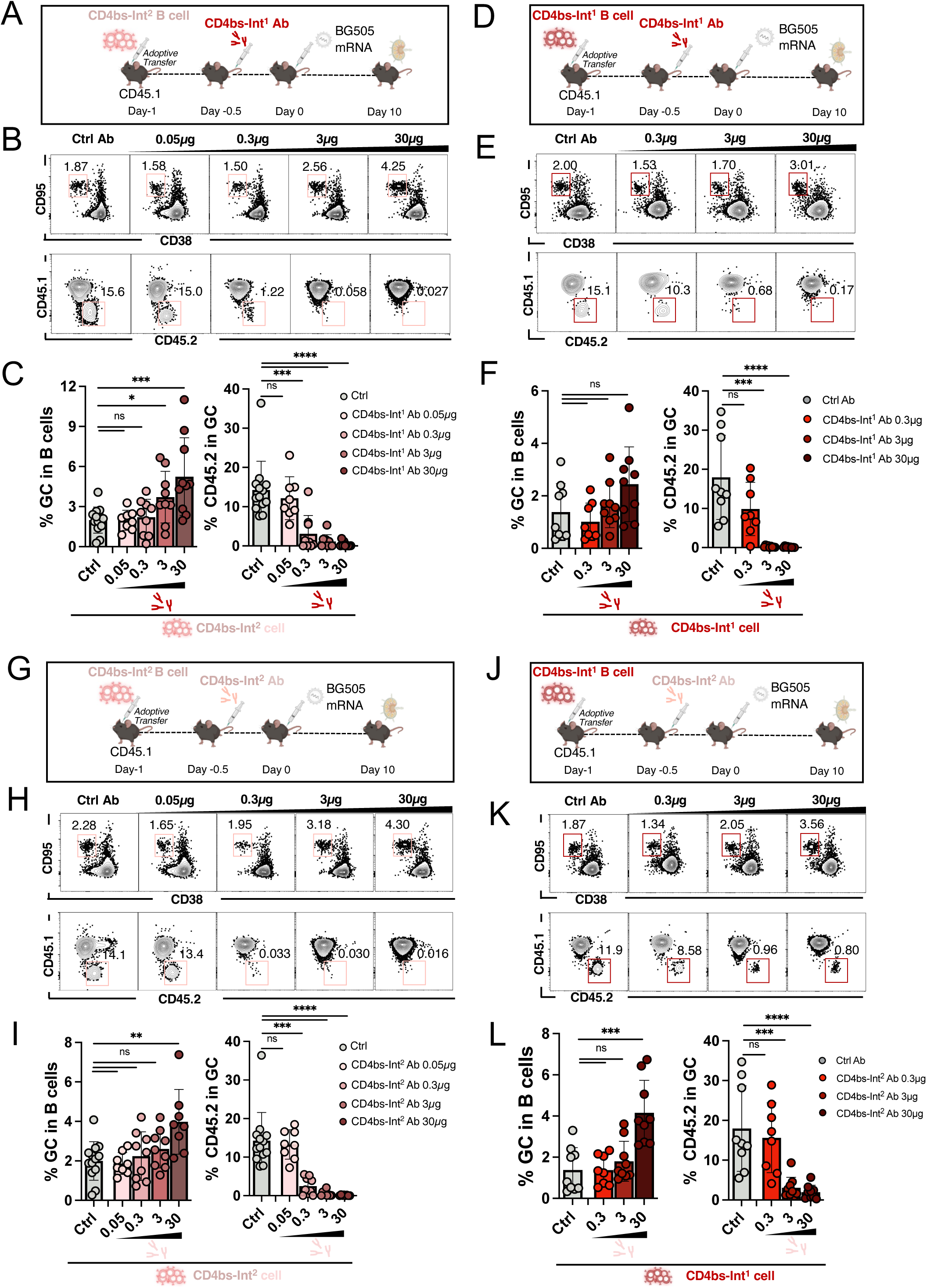
Affinities and concentrations of antibodies competing for the same epitope determine GC response. (A) Schematic of CD4bs-Int^1^ Ab blocking experiments for CD4bs-Int^2^ KI B cells. (B) (upper) Representative FACS plots of GC CD4bs-Int^2^ CD45.2 B cells obtained from dLNs after CD4bs-Int^1^ Ab pre-injection and BG505 mRNA immunization. Events were pre-gated on lymphocytes/singlets/live/CD4^-^CD8^-^F4/80^-^Gr1^-^/B220^+^ B cells and represent GC in B cells. (lower) Representative FACS plots of CD45.2 B cells. Events were pre-gated on lymphocytes/singlets/live/CD4^-^CD8^-^F4/80^-^Gr1^-^/B220^+^/CD38^-^CD95^+^ B cells and represent CD4bs-Int^2^ CD45.2 cells in GC. (C) (left) GC B cells as a percentage of total B cells and (right) CD45.2^+^ CD4bs-Int^2^ KI cells as a percentage of GC B cells at day 10. (D) Schematic of CD4bs-Int^1^ Ab blocking experiments for CD4bs-Int^1^ KI B cells. (E) Representative FACS plots of GC CD4bs-Int^1^ CD45.2 B cells obtained from dLNs at day 10. Gated as in (B). (F) (left) GC B cells as a percentage of total B cells and (right) CD45.2^+^ CD4bs-Int^1^ KI cells as a percentage of GC B cells at day 10. (G) Schematic of CD4bs-Int^2^ Ab blocking experiments for CD4bs-Int^2^ KI B cells. (H) Representative FACS plots of GC CD4bs-Int^2^ KI B cells obtained from dLNs after CD4bs-Int^2^ Ab pre-injection. Gated as in (B). (I) (left) GC B cells as a percentage of total B cells and (right) CD45.2^+^ CD4bs-Int^2^ KI cells as a percentage of total GC B cells at day 10. Control group reproduced from C. (J) Schematic of CD4bs-Int^2^ Ab blocking experiments for CD4bs-Int^1^ KI B cells. (K) Representative FACS plots of GC CD4bs-Int^1^ KI B cells obtained from dLNs after CD4bs-Int^2^ Ab pre-injection. Gated as in (B). (L) (left) GC B cells as a percentage of total B cells and (right) CD45.2^+^ CD4bs-Int^1^ KI cells as a percentage of total GC B cells at day 10. Control group reproduced from F. Thirty µg of irrelevant flu MEDI8852 Ab was pre-injected to each mouse in all Ctrl Ab groups. Figures represent data pooled from two experiments with 3–5 mice per condition. Each dot represents one mouse. Bars are mean + SD. Adjusted p-values (q-values) calculated using Kruskal-Wallis test followed by pairwise comparisons with BKY correction. Adjusted p-values (q-values) are indicated as follows: *p < 0.05, **p < 0.01, ***p < 0.001, ****p < 0.0001; ns = not significant.

We then repeated both adoptive transfers with the lower affinity CD4bs-Int^2^ mAb (*K*_D_, 46 nM for BG505) (**Fig. 6G&J**). As before, high mAb doses increased GC size. While 0.05 µg had no effect, CD4bs-Int^2^ mAb inhibited CD4bs-Int^2^ B cells in GCs beginning at a dose of 0.3 µg and reached near-total inhibition at 3 µg and 30 µg. (**Fig. 6H–I**). In contrast, while high mAb doses also increased GC size, 0.3 µg of CD4bs-Int^2^ mAb did not produce any significant decrease in CD4bs-Int^1^ B cells in GCs. Higher doses of CD4bs-Int^2^ Ab did reduce CD4bs-Int^1^ B cell numbers in GCs, however (**Fig. 6K–L**). Thus, antibody-mediated epitope masking suppresses GC participation in an affinity- and concentration-dependent manner, with feedback sensitivity governed by BCR affinity.

### Local plasma cells provide antibody feedback to GCs

To determine whether emerging plasma cells and subsequent antibody production could contribute to the differing GC kinetics observed for CD4bs-Int^1^ and CD4bs-Int^2^, we next interrogated plasma cell populations in and around GCs. GC formation occurs 5–7 days post-immunization and short-lived plasma cells are generated relatively early in the response^60^. As the frequency of CD4bs-Int^1^ B cells in GCs declined dramatically between days 7 and 16 post immunization, we focused on days 6 to 9 to capture this transition. Mice were adoptively transferred with either CD4bs-Int^1^ or CD4bs-Int^2^ cells and then immunized with 8 µg BG505 mRNA, after which flow cytometry was used to evaluate both CD45.2 recruitment to GCs and differentiation into plasma-like (CD138^+^) cells in and out of GCs in dLNs (**Fig. 7A**). At days 6 to 8, significantly more CD45.2^+^ KI cells were recruited to GCs after CD4bs-Int^1^ transfer compared to CD4bs-Int^2^-transfer, but the difference was no longer apparent by day 9 (**Fig. 7B–C**). Within GCs (inGC) at day 6, numbers of plasma-like (CD138^+^) cells were similar for CD4bs-Int^1^ (25%) and CD4bs-Int^2^ (18%). At days 7 and 8, CD138^+^ CD45.2 B cells were significantly higher in CD4bs-Int^1^ recipients (D7=24%; D8=15%) than CD4bs-Int^2^ recipients (D7=10%; D8=1%); but by day 9, few or no cells remained in GCs in either model (CD4bs-Int^1^=3.6%; CD4bs-Int^2^=0%) (**Fig.7B–C**). In unimmunized control mice, dLN from mice adoptively transferred with CD4bs-Int^1^ or CD4bs-Int^2^ showed no detectable CD138^+^CD45.2 cells in GCs from D6–D9 (**Fig. S8B**). In immunized mice, for local draining lymph node B cells not in GCs (exGC), the two lines were similar at day 6 and highly divergent after that: CD4bs-Int^1^ maintained ∼40% of plasma-like CD138^+^ cells exGC throughout; CD4bs-Int^2^ CD138^+^ cells exGC decreased from 25% at day 6 to 6.5% at day 9 (**Fig.7B–C)**. The exGC environment thus contained detectable plasma-like cells from day 6 to 9 for either line, but CD4bs-Int^1^-derived plasma cells remained at far higher percentages. This group of plasma-like CD138^+^ cells were also observed at day 8 inGC (9%) and exGC (30%) in CD4bs-Int^1^ recipient mice but not CD4bs-Int^2^ recipients after lower affinity DU172 protein immunization; the population decreased to below 1% between day 8 and 16 (**Fig. S7A&B**). The lower numbers of early CD138^+^ cells in dLN, especially inGC, could be related to longer GC residence in lower affinity DU172 immunization.

**Figure 7.**
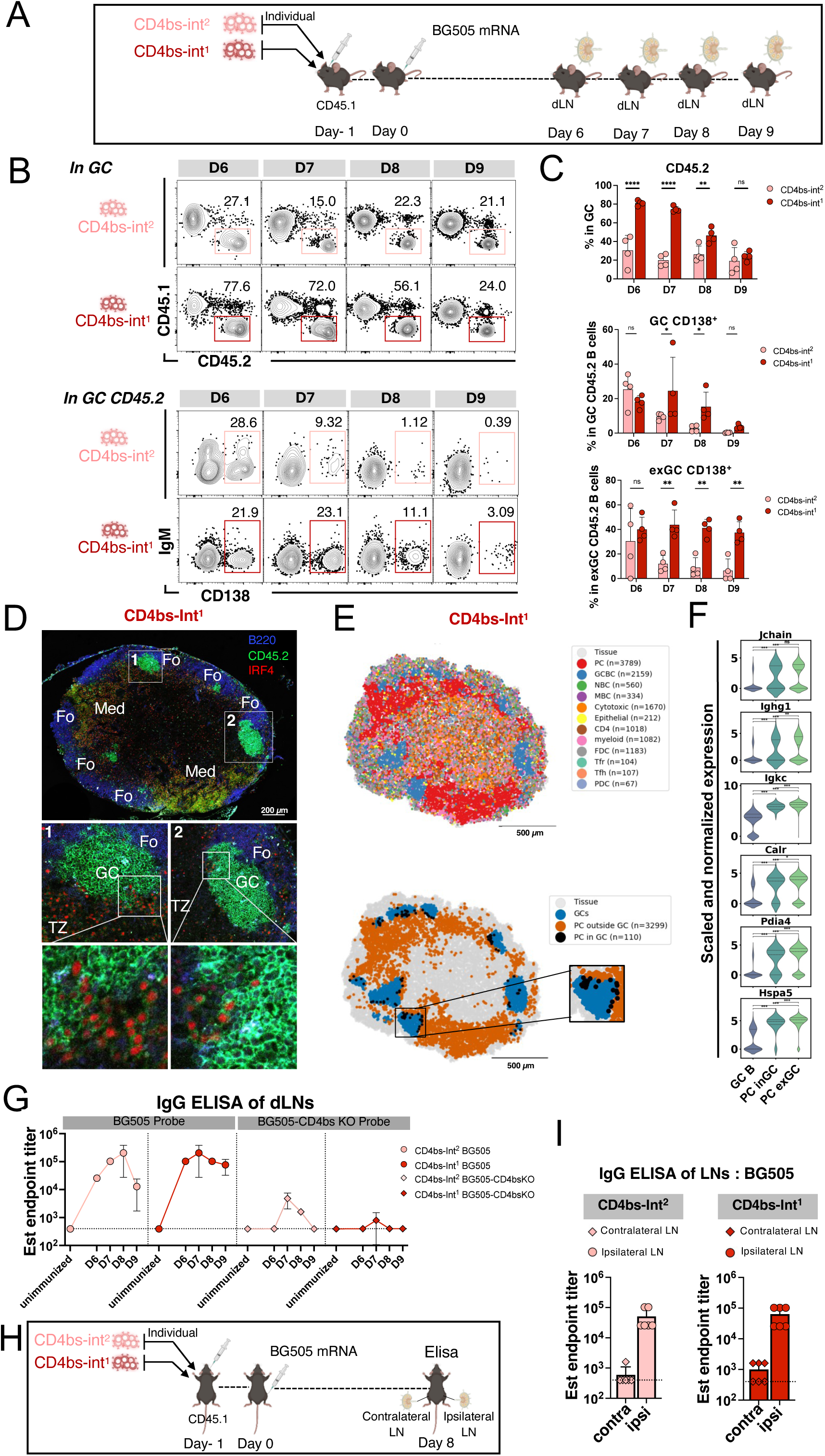
Plasma cells in and adjacent to GCs produce a local Ab pool that adjusts the GC response. (A) Schematic of plasma cell experiments. Recipient mice were adoptively transferred with either CD4bs-Int^1^ or CD4bs-Int^2^ KI B cells to reach a frequency of 50 in 10^6^ B cells before immunization with 8 µg of BG505 mRNA. Mice were sacrificed at days 6, 7, 8, and 9. (B) (upper) Representative FACS plots of CD45.2 KI B cells. Events were pre-gated on lymphocytes/singlets/live/CD4^-^CD8^-^F4/80^-^Gr1^-^/B220^+^/CD38^-^CD95^+^ B cells and represent CD45.2 KI cells in GCs. (lower) Representative FACS plots of plasma like cells. Events were pre-gated on lymphocytes/singlets/live/CD4^-^CD8^-^F4/80^-^Gr1^-^/B220^+^/CD38^-^CD95^+^/CD45.1^-^CD45.2^+^ B cells and represent plasma-like cells among GC CD45.2 B cells. (C) (upper) CD45.2^+^ KI cells as a percentage of GC B cells, (middle) CD138^+^ KI cells as a percentage of GC CD45.2 B cells, and (bottom) CD138^+^ KI cells as a percentage of exGC CD45.2 B cells at days 6, 7, 8, 9. Dots are individual mice, bars are mean + SD. Adjusted p-values (q-values) calculated by 2-way ANOVA with pairwise post-hoc comparisons adjusted using BYK correction. Adjusted p-values (q-values) are indicated as follows: *p < 0.05, **p < 0.01, ***p < 0.001, ****p < 0.0001; ns = not significant. (D) Representative immunofluorescent staining of dLN isolated from mice transferred with CD4bs-Int^1^ B cells at day 6 post-immunization by BG505, staining with B220 (blue, surface), CD45.2 (green, surface), and IRF4 (red, nucleus). Large tile presents overview of architecture of the whole LN, inset boxes present individual GCs and subsets of PCs in GC. Fo, Follicle; MC, Medullar cords; TZ, T cell zone. Scale bar: 200 µm. (E) Slide-seq spatial map of dLN isolated from mice with CD4bs-Int^1^ B cells 6 days post-immunization by BG505. Mapping of dLN is (upper) colored by cell type annotations from gene expression profiles, or (lower) colored by anatomical location inGC or exGC. Plasma cells (PC); GC B cells (GCBC); naïve B cells (NBC); memory B cells (MBC); cytotoxic T cells (Cytotoxic); epithelial cell (Epithelial); CD4 T cells (CD4); follicular dendritic cells (FDC); follicular regulatory T cells (Tfr); follicular helper T cells (Tfh); plasmacytoid dendritic cells (PDC). n = number of subset cells. Non-serial sections of the same dLN were used as in (D). Scale bar: 500 µm. (F) Violin plots of expression level of genes related to Ab secretion in GC B cells and PCs in and out of GC by anatomical location in the dLN at day 6. Pairwise group comparisons were performed using two-sided Mann–Whitney U tests, followed by Holm–Bonferroni correction for multiple comparisons. Adjusted p-values (q-values) are indicated as follows: *p < 0.05, **p < 0.01, ***p < 0.001, ****p < 0.0001; ns = not significant. (G) ELISA quantification of BG505-binding (circle) and BG505-CD4bs-KO-binding (diamond) IgG from 100 µl of dLN homogenates of CD4bs-In^1^(red) or CD4bs-Int^2^ (pink) recipient mice immunized with BG505 mRNA or left unimmunized. A 5x dilution multiplier was applied to obtain the final estimated (est) endpoint titer of dLNs. Dots represent mean values of technical triplicate from homogenates generated from one popliteal LN from each of the four mice pooled at days 7, 8, and 9. Unimmunized dots include all data collected at days 7, 8, and 9 in groups left unimmunized after corresponding adoptive transfers. Dotted lines represent LOD. (H) Schematic of single-side immunization experiment for CD4bs-Int^1^ and CD4bs-Int^2^ KI B cells. Recipient mice were adoptively transferred with CD4bs-Int^1^ or CD4bs-Int^2^ KI B cells respectively to reach a frequency of 50 in 10^6^ B cells before immunization with 8 µg of BG505 mRNA. Mice were sacrificed at day 8 for ipsilateral dLNs and contralateral LNs as controls. (I) ELISA quantification of BG505-binding IgG from 100 µl of dLN homogenates ipsilateral dLNs (circle) and contralateral LNs (diamond) of CD4bs-In^1^(red) or CD4bs-Int^2^ (pink) recipient mice immunized with BG505 mRNA or left unimmunized. A 5x dilution multiplier was applied to obtain the final est endpoint titer of dLNs. For each group, homogenates were generated from one popliteal LN from each of the four mice pooled into 100 µl of buffer. Technical triplicates were performed for each group. Dots represent individual values pooled from two experiments. Bars are mean ± SD. Dotted lines represent limit of detection.

The observation of CD138^+^ cells both inGC and exGC by flow cytometry pushed us to further explore the spatial distribution of plasma-like cells in the dLN. To determine localization, we first deployed immuno-fluorescence (IF) staining in immunized mice adoptively transferred with either CD4bs-Int^1^ (**Fig. 7D**) or CD4bs-Int^2^ (**Fig. S8C**); dLN from mice adoptively transferred with CD4bs-Int^2^ but left unimmunized were also examined as a control (**Fig. S8D**). Nucleus staining for IRF4 was utilized for better visualization of plasma-like cell colocalization. Multiple GCs formed by 6 days after BG505 immunization in the B cell follicles at the cortex of the dLN in adoptively transferred mice (**Fig. 7D & S8C**). Some GCs were comprised almost entirely of CD45.2 KI cells. Of those CD45.2 KI cells within GCs, some were also positive for IRF4 nucleus staining, indicating that they were plasma-like; CD45.2 plasma-like (IRF4^+^) cells were primarily observed in small groups at GC borders (**Fig. 7D, Fig. S8C**). Using Slide-seq^61,62^, we then developed a clearer and more comprehensive whole-distribution map of plasma-like cells in dLN. Plasma-like cells, assigned by Robust Cell Type Decomposition (RCTD)^63^ using an immune scRNA-seq reference dataset^64^, were observed both within and around GCs, accumulating in the medullary cords, paracortex, or T-B border (**Fig. 7D–E; Fig. S8C, S8E–F**). Quantification of sufficiently sized GCs (>=100 beads) showed an average of 1.5 PCs per 100 GC cells in the inner GC core, 6.6 per 100 in the outer GC ring, and 20.3 per 100 in the 30 µm periphery of the GC (**Fig. S8G-H**). Gene expression analysis found upregulation of Ab secretion-related genes, including secretion components (*Jchain*, *Ighg1*, *Ighg2b*, and *Ighg2c*), as well as protein folding machinery (*Calr*, *Pdia4*, and *Hspa5*), in PCs within the anatomical range of the GC relative to non-PC GC B cells; expression by inGC PCs reached similar or slightly lower levels compared to PCs exterior to the GC (**Fig. 7F, Fig. S8I**), suggesting potential Ab secretion functions in those cells.

To determine whether these plasma cell populations in and around the GC produced Ab, dLNs from different time points were mechanically disrupted in buffer and ELISA against BG505 was performed on the resulting supernatant. Based on prior measurements of murine LN volumes^65–70^, we estimated that a minimum of a 5x dilution occurs at the mechanical disruption stage; we therefore applied a 5x correction to estimate endpoint titer in the dLNs. Compared to unimmunized mice adoptively transferred with the same KI cells, high titers of IgG were detected in dLNs at 6 to 9 days after immunization. For CD4bs-Int^2^, the estimated endpoint titer for dLN IgG increased from 3×10^4^ at day 6, to 1×10^5^ at day 7, to a peak of 2×10^5^ at day 8, and then dropped to 1×10^4^ at day 9. High-affinity CD4bs-Int^1^ presented a higher and more stable dLN IgG titer curve above 8×10^4^ from day 6 to 9, with a slightly earlier peak of 2×10^5^ at day 7. At day 7, a small BG505-CD4bs-KO probe peak was observed for CD4bs-Int^2^ (at 5×10^3^) and CD4bs-Int^1^ (8×10^2^), potentially indicating a small amount of IgG produced by host mouse B cells targeting other epitopes (**Fig. 7G)**. Compared to IgG, IgM titers peaked at a lower level (5×10^3^) and early (by day 6) in dLNs of both types of transfer recipients, with low affinity CD4bs-Int^2^ persisting until day 9 while, in contrast, high affinity CD4bs-Int^1^ dropped at day 7. No IgM were detected above the detection limit of 250 for either the CD4bs-Int^1^- or CD4bs-Int^2^-transferred unimmunized group, or by BG505-CD4bs-KO probes (**Fig. S8J)**. The observation that neither IgG nor IgM were detected in the dLN of unimmunized adoptive transfer recipients excludes the possibility of a base from self-reactivity (**Fig. 7G, Fig. S8J**). Local dLN titers may be higher than those in circulation at early timepoints (**Fig. S8K**). To more precisely establish the localization of Ab production to the dLN, we performed single-side immunization on recipient mice and compared LN homogenate from ipsilateral dLNs and the contralateral LNs (**Fig. 7H**, **Fig. 7I, Fig. S8L**). At day 8, the ipsilateral dLNs showed markedly higher average IgG titers in both CD4bs-Int^2^ recipient (5.1 x 10^4^) and CD4bs-Int^1^ recipient (6.4 x 10^4^), while the contralateral LNs showed much lower IgG titers just above LOD (6 x 10^2^ for CD4bs-Int^2^; 1 x 10^3^ for CD4bs-Int^1^) (**Fig. 7I, Fig. S8L**). This result suggests that early Ab feedback could be more local than systematic.

To determine the affinities of the early antibody pool, mutated BCR sequences from day 7 GC B cells were expressed as IgG mAbs in vitro and tested for affinity against the BG505 trimer. In day 7 GC, mAbs isolated from CD4bs-Int^1^-transfer recipients had relatively high affinities, with *K*_D_ values ranging from 3 nM to 19 nM and a median *K*_D_ of 5 nM, while mAbs from CD4bs-Int^2^ recipients had relatively lower affinities, with *K*_D_ values ranging from from 34 nM to 2.5 µM and a median *K*_D_ of 85 nM (**Fig. S8M**). This indicated a maintenance of an at least 10-fold overall affinity gap between the Ab reservoirs generated by these two lineages. In sum, high early plasma-cell and antibody abundance in dLNs demonstrate the existence of a local antibody feedback loop that rapidly tunes GC competition.

## Discussion

The classical model of germinal center (GC) selection is grounded in Darwinian principles, where higher-affinity B cells progressively outcompete lower-affinity clones for antigen and survival signals^71^. Our data reveal an additional layer of regulation in which antibody feedback shapes GC dynamics in a nuanced manner. Using mouse models with BCRs of defined affinities and epitope specificities to the same immunogen, we found substantial initial recruitment of both high- and low-affinity B cells, but only lower affinity cells remained in GCs over time. This persistence was not a fixed characteristic of particular B cell lines but rather depended on their affinity for the presented immunogen. Importantly, B cells targeting the same epitope with similar affinities could be costimulated without changing their individual GC residency patterns, whereas high-affinity clones suppressed lower-affinity counterparts targeting the same epitope. Spatial transcriptomics revealed the presence of plasma-like cells near GCs as a probable source of local secreted antibody feedback mediating this suppression. Together, these findings suggest that in addition to Darwinian affinity selection, antibody feedback may provide a critical regulatory “brake,” enforcing affinity thresholds that shape B cell selection and exit from GCs, while also promoting epitope spreading as lower-affinity clones to other epitopes are not suppressed. This dual mechanism has significant implications for vaccine strategies aiming to balance breadth and potency by modulating affinity and feedback dynamics.

B cells must meet a minimum affinity-to-antigen “floor” to enter GCs and undergo further affinity maturation^3,5,11,12,72–74^. With increasing affinity, diminished selective pressure on antigen–BCR binding may occur^3,18,19^; high affinity variants may emerge, but a ceiling exists above which greater affinity maturation does not provide a selective advantage over other high affinity cells. B cells with higher initial BCR affinities to antigen have been observed in the classic hapten-based system to accumulate fewer V_H_ mutations than their low-affinity counterparts despite similar mutation rates, due to either a relaxation in positive selection or to negative selection as the likelihood increases for high affinity BCRs that any mutation diminishes affinity^13^. Similarly, in our real-world HIV-trimer-based system, the presence of high affinity competitors seemed to drive increased SHM rates in some high affinity B cells, potentially due to stronger selection. Antibodies produced by those first activated B cells in GC can modulate this selective process by directly competing for antigen or inducing apoptosis in lower-affinity B cells^25^; high-affinity antibodies may be induced quite early, without affinity maturation^19^. Recent studies have highlighted the blocking effect injected high affinity Abs can exert on immune responses^27,28^. In agreement with these studies, our findings suggest an Ab-driven self-modulation loop acts as a “brake”, reinforcing the upper affinity maturation ceiling in GCs. Notably, we found this brake to be an intra-epitope phenomenon reproducible by passive antibody administration, suggesting that epitope masking by early antibody production may be the source in this system. The epitope-specific nature of the brake is congruent with recent findings in an influenza infection model, where affinities from interclonal plasma cells (PC) differed by factors of a thousand or more, while intraclonal PCs only differed by factors of 10 to 30^17^. This intraclonal ceiling effect produced by Ab braking may drive B cell diversity in GCs and subdominant responses observed by multiple studies^28–30,75,76^. Interestingly, feedback from serum Abs has been observed not only hindering but also enhancing responses^27–29,77^. The increase in total GC sizes after high doses of Ab in our passive transfer experiments is suggestive of this latter phenomenon, and is notable in light of recent work suggesting that Abs can essentially adjuvant mRNA-LNP immunization^28^. In contrast to the “brake”, Ab-driven self-modulation loop may also serve as the intra-epitope “filter” to exclude the low-affinity lines from GC entry or residence and ensure only the “best in class” lines are favored—though avidity effects may in some cases rescue low affinity B cells from disadvantages in GC participation^78^. Together, the combination of initial BCR affinity to antigen and the timing of high affinity antibody secretion may establish an Ab-driven self-modulating loop, in which the “brake” drives diversity while the “filter” drives potency.

Much of our understanding of Ab feedback describes the relationship between the humoral response and serum Ab titers and affinities^28,79,76,27^, a mechanism particularly relevant to immune recall in a landscape of pre-existing circulating Abs. In contrast, our observations of early Ab titer in dLN and PC in and around the GC suggests that a local, real-time PC-driven feedback loop determines B cell composition in primary GCs. Plasma fate commitment occurs in the light zone (LZ) with the upregulation of *Irf4*^16,80–82^. Immunofluorescent staining using single markers, such as IRF4, BLIMP1, CD138, or cytosol IgG1, had placed PC precursors predominantly in the T-B border, with rare subsets in the GC^80,81,83,84^. Gene expression analysis of plasma-like cells (IRF4^+^) defined as GC-resident on the basis of CD38^-^Fas^+^ are similar to the total plasma-like population (CD138^+^TACI^-^)^85^. Our Slide-seq data provide a clearer picture of PC distribution in dLN and confirms the existence of inGC PCs designated by a comprehensive gene signature. Similar levels of upregulation of secretion components and protein folding machinery in inGC PCs and exGC PCs implies an Ab secretion function in those cells; recent observations of unusually low extracellular protease levels inside B cell follicles in mouse LN^86^ further support the possibility of a local Ab pool. Like the GC response itself, post-GC PC proliferation is affinity-dependent^87^, and our own observations suggest more inGC PCs from higher-affinity lines were present throughout early GC development. Thus, in addition to precursor frequency, Ag affinity, and Ag avidity^88^, a localized PC population may serve as the source of a local high affinity secreted Ab pool determining the outcome of B cell competition within GCs.

Affinity- and kinetics-driven competition and Ab feedback have substantial implications for the vaccine-driven development of breadth, particularly in contexts where the maturation of multiple B cell lineages are driven by the same immunogens to achieve a broadly neutralizing response, such as HIV sequential immunization^32,39,41,89–91^. While great progress has been made in simultaneously priming diverse bnAb precursors^92,93^, little is known about how crosstalk between minimally matured antibodies and B cells will impact late-stage boosting strategies. The lack of inter-epitope B cell competition in our assays indicates that the brake on intra-epitope B cell development may be essential to a multi-epitope response. Where long GC reactions are preferable, the use of a lower-affinity immunogens may be required to release the self-braking mechanism. Furthermore, affinity of an antigen to the broader antibody and B cell repertoire must be considered. Though not ascribed to antibody production, the presence of higher affinity non-precursor same-epitope responses has previously been observed as a limiting factor in B cell GC residence^12^. BnAb development strategies involving sequential immunization should aim to identify functional affinity ranges for each immunization stage. A related concept, that the “affinity drop” between sequential antigens should not be too high or too low, was previously described, though also not ascribed to antibody production^39^.

In sum, our study elucidated a multi-faceted Ab feedback loop in B cell competition for the same immunogen, which served as a self-modulating affinity-dependent “brake” for development of B cell lineages to the same epitope, as well as a “filter” excluding lower affinity B cells, while preserving a parallel evolutionary path for lineages to other epitopes. The use of vaccine models applicable to pre-clinical development provides direct applications to fine-tuning vaccine strategies to avoid inhibitory antibody feedback.

### Limitations of the Study

Our findings demonstrate that inherent affinity significantly impacts the durability and magnitude of GC B cell responses. In terms of tools used here, while we found antibody feedback to be compelling based on multiple lines of evidence in our data, more direct tests, potentially using antibody-production-deficit B cells, such as Blimp-1 knock outs, remain for future investigations. Additionally, Slide-seq, while a powerful approach, has a lower capture rate than traditional RNA sequencing. Furthermore, in terms of our overall model system, where cross-epitope comparisons are made, we must note that the bnAbs used in our BCR knock-in mice differ in binding stoichiometry (monovalent vs trivalent), binding modality (HCDR3 vs V_H_-gene dominant), approach geometry, and epitope composition. Antigen-BCR binding stoichiometry and approach geometry may influence the degree of BCR crosslinking on antigen-decorated surfaces, while binding modality and epitope composition could affect BCR clustering. In particular, signaling thresholds may differ for BCRs recognizing predominantly proteinaceous epitopes to those engaging carbohydrate-rich epitopes, as glycan-mediated interactions may exhibit distinct binding kinetics or activation dampening via CD22 interactions^94^. Antigen presentation may also complicate a purely affinity-driven picture, as the membrane-anchored presentation of the immunogen used here may be subject to quite different modulating forces than a soluble immunogen^57,58,95^, though no obvious differences in individual kinetics, at least, were observed in our soluble protein experiments. Nevertheless, consistent affinity-dependent effects were observed both when varying BCR affinity through diverse bnAb knock-ins and when varying antigen affinity through different immunogens.

## Supporting information

Supplemental Figures 1 through 8

## Resource Availability

Model animals and antibodies minBG18.6-1, minBG18.6-2, minBG18.6-3, minBG18.6-4, minBG18.11-1, minBG18.11-2, minBG18.11-3, minBG18.11-4, N6-I3-D7-1, N6-I3-D7-2, N6-I3-D7-3, N6-I3-D7-4, N6-I3-D7-5, N6-I3-D7-6, N6-I3-D7-7, N6-I3-D7-8, N6-I3-D7-9, Min12A21-D7-1, Min12A21-D7-2, Min12A21-D7-3, Min12A21-D7-4, Min12A21-D7-5, Min12A21-D7-6, Min12A21-D7-7, Min12A21-D7-8, Min12A21-D7-9 are available from corresponding author (FDB) on request, under a standard material transfer agreement with Massachusetts General Hospital. Slide-seq data has been deposited on Broad Institute Single cell portal, and the repository URL is listed in the Key Resources Table (https://singlecell.broadinstitute.org/single_cell/). Custom code used for analysis has been deposited on GitHub, and the repository URL is listed in the Key Resources Table (https://github.com/). BCR sequences will be available upon publication in GenBank, and Key Resources Table will be updated to contain accession numbers. The code and sequence data are available as of the date of publication. Plasmids or recombinant proteins for immunogens and sort reagents related to BG505 MD39.3 and DU172 MD39.2; or antibodies 12A21, min12A21, N6-I2, N6-I3, N6, PCT64-18D; or SPR reagents in this study, are available from WRS under a material transfer agreement with the Scripps Research Institute. mRNA-LNP vaccine construct for BG505 MD39.3 can be made available from SH if the recipient and Moderna are able to agree upon the terms of a material transfer agreement.

## Acknowledgements

We would like to thank all members of the Batista lab for experimental help, as well as the Ragon Flow Cytometry Core and Scientific Editing Platform. We would also like to thank the Irvine lab (Scripps) for the provision of SMNP. Funding was provided by the Gates Foundation Collaboration for AIDS Vaccine Discovery (CAVD) grants INV009585 and INV046626 (to FDB); and NAC INV-007522, INV-008813, and INV-034657 (to WRS); National Institute of Allergy and Infectious Diseases (NIAID) UM1 AI144462 (Scripps Consortium for HIV/AIDS Vaccine Development) (to WRS and FDB); the IAVI Neutralizing Antibody Center (NAC) to WRS, and flexible funding from the Ragon Institute of Mass General Brigham, MIT and Harvard (to FDB).

## Author Contributions

YY designed, planned, and performed experiments, analyzed and interpreted the data, and wrote the paper. XW contributed to the design and conduct of experiments. ZX coordinated sample processing and cell sorting. DLVB designed immunogens and antibodies, performed SPR analysis, and edited the paper. RHL performed and analyzed section staining and Slide-seq. KMM and CAC designed antibodies and probes. JMSteichen designed immunogens and antibodies. LX assisted with confocal and in vitro stimulation experiments, and mouse colony management. PMV assisted with peritoneal lavage and FACS recording. MA and JHK provided antibodies. JMShen reviewed the paper. AV performed section staining and Slide-seq. OK performed SPR. JDA performed glycan profiling. AAA assisted in sorting experiments. AA assisted in sample processing. BC assisted with spatial transcriptomic analyses. EG, NA, NP, and RT purified immunogens and probes. JREP and QAP generated mouse lines. HN assisted in mice genotyping and breeding. AE assisted in affinity experiments. BK performed 10x sequencing. MK expressed immunogens and mAbs, purified mAbs. AL performed SPR. TP assisted in animal experiments. DL expressed immunogens and mAbs. SE expressed immunogens and mAbs. XL managed mouse colonies and lab resources. JEW performed and analyzed 10x sequencing experiments. SRW assisted in manuscript drafting and editing. SH produced mRNA-LNP vaccine constructs. MC performed glycan profiling. UN oversaw and performed mouse line generation. SL, WRS, and FDB conceived of and oversaw projects.

## Declaration of generative AI and AI-assisted technologies in the writing process

During the preparation of this work the authors used ChatGPT in order to edit and improve the readability of some text. After using this tool, the authors reviewed and edited the content as needed and take full responsibility for the content of the published article.

## Supplemental Information

Document S1. Figures S1–S8

## STAR Methods

### Experimental model and subject details

Η^N6-I3/N6-I3^κ^N6-I3/N6-I3^ (CD4bs-Int^1^), Η^min12A21/min12A21^κ^min12A21/min12A21^ (CD4bs-Int^2^), Η^N6-I2/N6-I2κN6-I2/ N6-I2^ (CD4bs-Int^3^), Η^PCT64-18D/PCT64-18D^κ^PCT64-18D/PCT64-18D^ (V2-Int^1^), Η^minBG18.6/minBG18.6^κ^WT/WT^ (V3-Int^1^), and Η^minBG18.11/minBG18.11^κ^WT/WT^ (V3-Int^2^) BCR knock-in (KI) mouse lines were generated on the background of C57BL/6J (CD45.2^+/+^) mice, as described previously^55,56^. Note, the HC sequences for V3-Int^1^ and V3-Int^2^ differ from the previously published minBG18.6 and minBG18.11 HC sequences by one amino acid in the J_H_ gene (ARNAIRIYGVVALGEWFHYGMDVWGQGTAVTVSS for V3-Int^1/2^; ARNAIRIYGVVALGEWFHYGMDVWGQGT<u>T</u>VTVSS for minBG18.6/11)^40^. All transgenic mouse lines were generated in the animal facility of the Gene Modification Facility (Harvard University). Subsequent breeding, colony maintenance, and experimental procedures were performed at the animal facility of the Ragon Institute of Mass General Brigham, MIT, and Harvard. For experiments, wild-type (WT) adult male B6.SJL-Ptprca Pepcb/BoyJ (CD45.1^+/+^) mice between the age of 8–12 weeks were purchased from The Jackson Laboratory (Bar Harbor, ME). Experimental mice were housed at the animal facility of the Ragon Institute with free access to food and water, controlled temperature, and a 12:12 hour light-dark cycle. All animal experiments were conducted in accordance with the Institutional Animal Care and Use Committee (IACUC) of Massachusetts General Hospital (MGH)’s approved Animal Study Protocols 2016N000286 and 2016N000022. The MGH Center for Comparative Medicine (CCM) is an Association for Assessment and Accreditation of Laboratory Animal Care (AAALAC) International-approved program.

### Immunogen and probe design

In this work we sought to employ native-like immunogens derived from wild-type HIV-1 isolates to present unmodified epitopes capable of engaging moderately evolved broadly neutralizing antibodies (bnAbs) targeting diverse neutralizing supersites on Env: CD4-binding site (CD4bs), V2-apex (V2), and V3-glycan (V3). We selected Env from two strains whose soluble antigens fulfill two independent criteria to evaluate how affinity differences on both the antigen and BCR sides influence antibody feedback mechanisms: (1) Env-1 must show detectable binding affinity to the diverse minimally mutated bnAbs of interest (PCT64-18, minBG18.6, minBG18.11, min12A21 and N6-I3), and (2) Env-2 must display significantly lower affinity than Env-1 for CD4bs antibodies. BG505 (clade A) and DU172-17 (clade C) served as Env-1 and Env-2, respectively, fulfill these criteria (**Fig. S6B-C**). BG505 MD39.3 construct design and characterization was previously described^58^. DU127-17 was selected because it is resistant to neutralization by several CD4bs-targeting bnAbs and its recombinant antigens display low affinity against VRC01-class bnAbs^52,96^. DU172 MD39.2 was designed using established HIV-1 Env stabilization strategies: (1) SOSIP mutations for improved stability^97–99^, (2) C-terminal truncation at residue D664 for enhanced homogeneity of trimeric pre-fusion gp140^99,100^, (3) MD39 mutations for improved antigenic profile, expression yield, and thermostability^39^, and (4) replacement of the furin cleavage site with a non-cleavable linker termed link14 between gp120 and gp41 (linker sequence: SHSGSGGSGSGGHA)^40^, where we use the terminology MD39.2 to denote a cleavage-independent MD39 stabilized trimer (**Fig. S6A**)^58^. Mass spectrometry analysis and antigenic profiling against a panel of neutralizing and non-neutralizing antibodies performed by BLI confirmed DU172 MD39.2 constructs exhibit the expected N-linked glycan profile and adopt a trimeric pre-fusion conformation (**Fig. S6B-D**).

Probes for BG505 and DU172 were generated by addition of His-Avi-tag (HHHHHHGGSGGSGLNDIFEAQKIEWHE) and His-tag (HHHHHH) sequences at the C terminus of gp41 after residue D664. while His-tagged trimers served as ELISA probes, and His-Avi-tag SOSIP served as FACS probes.

His- and His-Avi-tagged epitope-specific KO reagents were generated by introducing four VRC01-class-specific KO mutations (280R, 365L, 368R, and 371R) onto BG505 MD39.3 and DU172 MD39.2 SOSIP trimers as previously described^89,101^. His-Avi-tagged V2-apex epitope-specific KO mutants were created by introducing R169E and K171E mutations that abrogate binding by long-HCDR3 Apex bnAbs and related precursors^95^. His-Avi-tagged V3-glycan epitope-specific KO variants were produced by introducing R327D, H330K and N332T mutations^40^. All HIV-1 Env residues are denoted using HxB2 numbering.

### Immunogen, probe, and Ab production

For protein immunogen production, untagged trimeric immunogens were produced by transient transfection of HEK-293F cells (ThermoFisher) and purified by gravity-flow affinity chromatography using *Galanthus nivalis* lectin resin (Vectorlabs) followed by SEC using Superdex 200 Increase 10/300 GL columns (Cytiva). The homogeneity and molecular weight of antigens was confirmed by size-exclusion chromatography-multi-angle light scattering (SEC-MALS) in PBS using Superdex 200 Increase 10/300 GL columns (Cytiva) with an isocratic flow of 0.5 mL/min followed by DAWN HELEOS II and Optilab T-rEX detectors (Wyatt Technology). Endotoxin levels in immunogen preparations were confirmed to be <5 EU/mg of endotoxin using an Endosafe nexgen-PTS instrument (Charles River).

For mRNA immunogen production, the amino acid sequence encoding BG505 MD39.3 (gp151) was provided to Moderna for production and formulation of mRNA-LNP immunogens.

For probe production, His-tagged and His-Avi-tagged trimeric antigens were produced by transient transfection of HEK 293F cells (ThermoFisher) and purified by immobilized metal ion affinity chromatography (IMAC) using HisTrap excel columns 5 mL (Cytiva) followed by size-exclusion chromatography (SEC) using Superdex 200 Increase 10/300 GL columns (Cytiva). The homogeneity and molecular weight of antigens was confirmed by size-exclusion chromatography-multi-angle light scattering (SEC-MALS) in PBS using Superdex 200 Increase 10/300 GL columns (Cytiva) with an isocratic flow of 0.5 mL/min followed by DAWN HELEOS II and Optilab T-rEX detectors (Wyatt Technology). Biotinylation of His-Avi-tagged trimers was performed using BirA (Avidity) and purified by SEC with Superdex 200 Increase 10/300 GL columns (Cytiva) to remove unconjugated biotin molecules.

For Ab production, sequences encoding the antibody Fv regions were synthesized by GenScript and cloned into antibody expression vectors pCW-CHIg-hG1 and pCW-CLIg-hk for heavy and light chain genes, respectively. Monoclonal antibodies (mAbs) were produced using transient transfection of HEK 293F cells (ThermoFisher) and purified by gravity-flow affinity chromatography using rProteinA Sepharose Fast Flow resin (Cytiva). Min12A21, N6-I3, minBG18.6, and minBG18.11 antibody mutants were produced by GenScript using the TurboCHO expression service. All antibodies were produced as human IgG1s.

SMNP was provided by the Irvine lab of Scripps Research Institute^59^.

### Bio-layer interferometry (BLI)

Native-like conformation of soluble DU172 gp140 antigens was confirmed by Bio-Layer Interferometry (BLI) using a panel of broadly neutralizing (quaternary-specific PGT151 and PGT145; V3-specific PGT121; CD4bs-specific N6 and min12A21) and non-neutralizing antibodies (CD4bs-specific F105 and B6; V3-specific 19b) and conducted on an Octet RED Instrument (FortéBio). Non-neutralizing antibodies bind non-native trimers and monomeric gp120; BG505 gp120 foldon trimer served as a negative control representing poorly assembled Envs (binding to non-nAbs and lack of binding to quaternary-specific bnAbs). Monoclonal antibodies were captured on anti-hIgG Fc capture (AHC) biosensors (Sartorius) at a concentration of 10 µg/mL in kinetics buffer (PBS, pH 7.4, 0.01% [w/v] BSA, and 0.002% [v/v] Tween 20) for 120 seconds after baseline determination. Association was measured for 120 seconds by dipping IgG-loaded biosensors into wells containing 1 µM recombinant SOSIP trimers. Dissociation was monitored for 120 seconds in kinetics buffer. Relative binding was determined by subtracting baseline absorbance and calculating the maximum response (nm) at endpoint of the association phase.

Biotinylated sort reagents of BG505 MD39.3, BG505 MD39.3 CD4bs-KO, DU172 MD39.2, and DU172 MD39.2 CD4bs-KO were loaded onto streptavidin (SA) biosensors (Sartorius) at 25 µg/mL in kinetics buffer for 120 seconds after baseline determination. After reacquiring a kinetics buffer baseline, biosensors were transferred to wells containing 1 µM monoclonal antibodies (neutralizing and non-neutralizing IgGs) and allowed to associate for 120 seconds. The biosensors were dipped into kinetics buffer alone to monitor dissociation for additional 120 seconds. Relative binding was determined by subtracting baseline absorbance values from end-point measurements.

### Site-specific glycan profiling

N-linked site-specific glycan profiling was conducted as previously described to determine the degree of glycan occupancy and the extent of glycan heterogeneity (proportion of complex vs oligomannose/hybrid glycan types)^102^.

### Surface Plasma Resonance (SPR)

We measured kinetics and affinity of antibody-antigen interactions on Carterra LSA using CMDP Sensor Chip (Carterra) and 1x HBS-EP^+^ pH 7.4 running buffer (20x stock from Teknova, Cat. No H8022) supplemented with BSA at 1 mg/mL. The chip surface for ligand capture was prepared following Carterra software instructions. About 800–1000 RU of capture antibody (SouthernBiotech Cat.no 2047-01) in 10 mM Sodium Acetate pH 4.5 was amine coupled, and regeneration buffer Phosphoric Acid 1.7% was injected three times per cycle for 60 seconds. Ligand concentrations of 1 µg/mL were used with a contact time of 5 minutes. Analyte samples (SOSIP trimers) were buffer exchanged into the running buffer using dialysis and analyte concentrations were quantified on NanoDrop 2000c Spectrophotometer (Thermo Fisher Scientific) using absorption signal at 280 nm. Raw sensograms were analyzed using Kinetics software (Carterra), interspot and blank double referencing, Langmuir model. We typically cover a broad range of affinities in our runs and the best referencing practices differ depending on how fast the dissociation rate is for a particular ligand. For fast dissociation rates (faster than 9e^-3^ 1/s) we use automated batch referencing that includes overlay y-align and higher analyte concentrations. For slow dissociation rates (9e^-3^ 1/s or less) we use manual process referencing that includes serial y-align and lower analyte concentrations. After automated data analysis by Kinetics software, we also performed additional filtering to remove datasets with highest response signals smaller than signals from negative controls. This additional filtering was performed automatically using an R-script (available upon request).

### B cell in vitro stimulation assay

Naive live B cells were purified from spleens of KI and WT mice to reach more than 90% purity using negative B-cell isolation (Miltenyi Biotec). 10^7^ per mL B cells were labeled with 2 μM CFSE (Thermo Fisher) for 5 min at 37°C then were washed with complete B cell medium [RPMI supplemented with 10% FCS, 25 mM Hepes, Glutamax, Non-Essential Amino Acids, penicillin streptomycin (Thermo Fisher), and 50 μM β-mercaptoethanol (MilliporeSigma)]. CFSE Labelled cells were then stimulated in complete B cell medium supplemented with combinations of 10 ng/mL IL4 (R and D systems), 10 ng/mL IL5 (R and D systems), and 50 ng/mL CD40L (R and D systems). After 3 days of culture, proliferation status was measure by percentages of CFSE^low^ cells in flow cytometry based on CFSE levels at day 0 for each well. Survival status was measure by percentages of DAPI^-^ cells after staining with DAPI (Thermo Fisher Scientific). The differentiation of plasmablasts was measured by percentages of CD138^+^ cells after CD138 (281-2) staining.

### Adoptive transfer and immunization

For adoptive transfer, spleens were collected from donor BCR transgenic mice with a C57BL/6J (CD45.2^+/+^) background. Spleens were crushed through a 70 µm cell strainer and subjected to pan B cell isolation kits (Miltenyi Biotec). Isolated B cells were then qualified for live cells on a LUNA-FX7 automated cell counter (Logos Biosystems) and adjusted to the desired number and volume (100 µl/mouse) in phosphate-buffered saline (PBS) before transfer to CD45.1^+/+^ recipient mice by intravenous (i.v.) injection through the orbital sinus, establishing frequencies of 20 in 10^6^ B cells except where otherwise specified. For mRNA immunization, mRNA-LNP immunogens were diluted to desired quantity and volume in PBS. Unless otherwise stated, diluted mRNA was injected at 2 µg per mouse in 100 µl per mouse, intramuscularly (i.m.) through gastrocnemius, 50 µl each leg. For protein immunization, 10 µg immunogens mixed with 5 µg of SMNP adjuvant were immunized to each mouse in 100 µl PBS subcutaneously (s.c.), 50 µl at each side of tail base.

### Flow cytometry and cell sorting

For single cell suspensions, inguinal, popliteal, and iliac lymph nodes were crushed through a 70 µm cell strainer, centrifuged and re-suspended in FACS buffer (2% fetal bovine serum/PBS). Single-cell suspensions were kept on ice after. Probes were conjugated with streptavidin-Alexa Fluor 488, streptavidin-Alexa Fluor 647, or streptavidin-Alexa Fluor 594 for at least 30 min. Cells were blocked by Fc block (clone 2.4G2, BD Biosciences) for 15 min and then pre-incubated with freshly conjugated specific probes for 15 min. Probe-stained cells were then co-incubated with surface antibodies for another 15 min. If sorting for 10x sequencing, cell barcodes (BioLegend) and anti-mouse CD45 hashtags (BioLegend) were incubated with cells for 15 min during the coincubation step. Cells were washed 3 times with FACS buffer and re-suspended with DAPI 1:5000 diluted in FACS buffer, after which they were loaded into a BD LSR Fortessa analyzer or a BD FACS Aria Fusion sorter. Sorted cells were kept on ice for subsequent 10x sequencing procedures or at -80° for Sanger sequencing procedures. Data analysis was performed using FlowJo software (TreeStar).

### Single-cell BCR sequencing

For 10x Genomics single-cell BCR sequencing, bulk sorted cells were loaded onto the 10x Genomics Chromium Controller at 5,000–20,000 cells per reaction and encapsulated in gel beads in emulsion. Single-cell gene expression, V(D)J, and hash-tag oligo libraries were constructed using the Next GEM Single-cell 5′ Reagent Kits v2 (10x Genomics, Pleasanton, CA) following the manufacturer’s protocol. Libraries were then quantified by Tapestation 4200 (Agilent, Santa Clara, CA) and the Qubit double-stranded DNA High Sensitivity assay (AAT Bioquest, Sunnyvale, CA). Qualified libraries were pooled and sequenced on the Nextseq2000 sequencer (Illumina, San Diego, CA). Finally, sequence data were analyzed using Cell Ranger pipelines and by customized analysis for specific KI sequences.

For experiments with fewer cell numbers, single cell 96-well plate PCR was performed, as described previously^12^.

### LN homogenization and Enzyme-linked immunosorbent assays (ELISA)

To test IgG or IgM specific end point titers in LN, a homogenizing preparation step is needed prior to ELISA: one popliteal LN from each of the four mice in each group was carefully isolated and washed in PBS three times before all four were pooled into 100 µl of PBS with 1x Complete EDTA-free protease inhibitor cocktail (Roche). Pooled dLNs were homogenized on ice and centrifuged at 300 g, 4°C for 5 min. Supernatants were centrifuged again at 14000 g, 4°C for 10 min. Supernatants were transferred to a clean tube as prepared dLN homogenate and stored at - 80°C for the next step.

Homogenate of dLN or mouse serum was then used for ELISA. Anti-His Ab (1 mg/ml, 50 µl per well) were incubated in 96-well high-absorption ELISA plates (NUNC/Corning) pre-coated with BG505 or epitope-specific-KO probes with His-tag (1 mg/ml, 50 µl per well) for 2 h at room temperature (RT). Plates were then washed 5 times with 0.05% Tween 20 in PBS (tPBS) and blocked with tPBS with 3% BSA for 2 h at RT. After washing, serial-diluted (2–5 folds depending on preliminary estimates of initial Ab titer) dLN homogenate or serum were incubated at 4°C for 4 h with requisite starting dilutions. Plates were washed again and incubated with 50 μl per well of Alkaline Phosphatase AffiniPure Goat Anti-Mouse IgG or IgM (Jackson Immuno Research) at 1:5000 in tPBS + 0.5 BSA FOR 1 h at RT. 50 μl per well of p-Nitrophenyl phosphate dissolved in ddH2O was added to each well for incubation of 20 min at RT for the final chromogenic reaction. Absorbance at 405 nm was determined with a Synergy Neo2 plate reader (BioTek). Endpoint titer was determined as dilution of the last serial-diluted well with an OD405 read over the threshold value, which was set as (mean + 3 x SD) of the OD405 read in all negative wells. A 5x dilution multiplier was applied to obtain the final est endpoint titer of dLNs.

### Immunofluorescence and histology

Frozen murine lymph node tissue sections were immersed in Harris hematoxylin for 15 seconds before rinsing in deionized (DI) water and washing in 1X PBS for 30 seconds. Sections were dipped in Scott’s Tap Water Substitute for 45 seconds and rinsed in DI water before they were dipped in 70% EtOH and 90% EtOH for 30 seconds each. Sections were immersed in alcoholic-eosin for 2 minutes and rinsed in DI water. Sections were dipped in 90% EtOH and 100% EtOH for 15 seconds each and dipped in 100% EtOH for 30 seconds. Sections were immersed in xylene for one minute, mounted, and coverslipped.

To verify the presence of adoptively transferred plasma cells in murine lymph node tissue sections, mounted 10 µm frozen tissue sections were fixed in 10% formalin for 10 minutes and washed three times in 1X PBS. Sections were permeabilized in 0.1% Triton X-100 for 15 minutes and washed three times in 1X PBS. Sections were stained with antibodies for adoptively transferred cells (Alexa Fluor 488 anti-mouse CD45.2, 104, Biolegend, 1:200), B cells (Alexa Fluor 594 anti-mouse/human CD45R/B220, RA3-6B2, Biolegend, 1:200), and plasma cells (Alexa Fluor 647 anti-IRF4, 3E4, Biolegend, 1:200) for one hour at room temperature. Sections were washed three times in 1X PBS, mounted, and coverslipped. Confocal imaging was performed on the TissueFAXS SL Q (TissueGnostics).

### Spatial transcriptomics: sample processing

Fresh frozen murine lymph node tissues were embedded in Tissue-Tek OCT compound and frozen on dry ice before transferring to -80°C for long-term storage. Frozen blocks were warmed to -20°C in a cryostat (Leica CM3050S) for 30 min prior to handling. Tissues were sliced at a 10 µm thickness and Slide-seq was performed on the fresh frozen sections as previously described^103^. Serial sections were taken for tissue staining at a later time. Spatial libraries were sequenced on an Illumina NextSeq 2000 P3 flow cell with the following read structure: 42 bases Read 1, 8 bases Index 1, 41 bases Read 2, 0 bases Index 2. Each library received approximately 250 million reads.

## Spatial transcriptomics: data analysis

As a quality control measure, we first removed beads with a unique molecular identifier (UMI) count of less than 100. Next, to delineate tissue boundaries and remove off-tissue beads, we performed Density Based Spatial Clustering of Applications with Noise (DBSCAN)^104^. To ensure tissue regions were continuous, we subsequently performed dilation, whereby beads initially removed by DBSCAN were added back to the tissue if they were located within a small radius of on-tissue beads.

Robust Cell Type Decomposition (RCTD)^63^ was used to assign cell types (plasma cell (PC), germinal center B cell (GCBC), cytotoxic T cell (Cytotoxic), follicular dendritic cell (FDC), myeloid cell (myeloid), CD4 T cell (CD4), naive B cell (NBC), memory B cell (MBC), epithelial cell (Epithelial), T follicular regulatory cell (Tfr), T follicular helper cell (Tfh), and plasmacytoid dendritic cell (PDC)) as singlets or doublets to each bead on Slide-seq data from lymph node samples. Mouse orthologs of human genes were identified using Ensembl BioMart^105^ based on reference scRNA-seq datasets^64^.

Germinal center regions were identified through marker gene expression (*Bcl6, Aicda, Rgs13*, and *Stnb1m)* and RCTD cell type assignments, including beads classified by RCTD as GCBC singlets or doublets containing GCBCs. These masks were merged, followed by DBSCAN clustering and spatial dilation.

To perform differential gene analysis, we used Cell-type Specific Inference of Differential Expression (C-SIDE)^106^ to compare plasma cells inside versus outside germinal centers.

### Statistical analysis

For Slide-seq data, analyses were conducted in Python 3.9.19 (The Python Software Foundation). For each gene/feature, pairwise group comparisons were performed using two-sided Mann–Whitney U tests. P-values were then adjusted for multiple testing using the Holm–Bonferroni correction.

All other statistical analyses were performed using Prism 10 (GraphPad). Normal distribution was not assumed. For comparisons involving two groups, Mann-Whitney’s test was utilized. For comparisons involving three or more groups, adjusted p-values (q values) were calculated by either 2-way Analysis of Variance (ANOVA) or nonparametric ANOVA (Kruskal-Wallis tests), as indicated in the legends; this was followed by post-hoc pairwise comparisons with Benjamini-Krieger-Yekutieli (BKY) correction for head-to-head comparisons.

Adjusted p-values (q-values) are indicated as follows: *p < 0.05, **p < 0.01, ***p < 0.001, ****p < 0.0001; ns = not significant.

