## Supplemental Figures 1 through 8 for "Antibody feedback establishes an affinity brake in the germinal center"

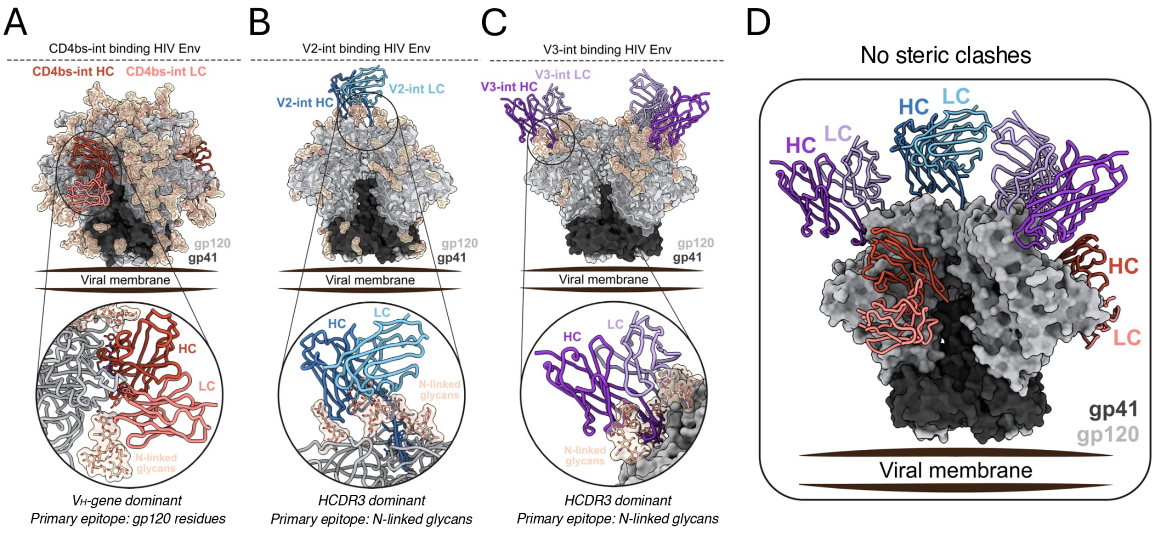


E

| Epitope | BCR knockin | bnAb | Key paratope | Binder type | Ab:Env ratio | Primary Epitope | HCDR3 length | $V_H$ %mut (aa) | $V_K/V_L$ %mut (aa) | Reference |
|---|---|---|---|---|---|---|---|---|---|---|
| CD4-binding site | CD4bs-int1 | N6-I3 | $V_H$-gene | $V_H$ binders | 3 | gp120 | 15 | 36 | 24 | Huang, 2016 |
| | CD4bs-int2 | min12A21 | $V_H$-gene | $V_H$ binders | 3 | gp120 | 15 | 17 | 8 | Jardine, 2016 |
| | CD4bs-int3 | N6-I2 | $V_H$-gene | $V_H$ binders | 3 | gp120 | 15 | 29 | 21 | Huang, 2016 |
| V3 glycan (N332) | V3-int1 | minBG18.6 | HCDR3 | Loop binders | 3 | N-linked glycans | 23 | 7 | 5 | Steichen, 2019 |
| | V3-int2 | minBG18.11 | HCDR3 | Loop binders | 3 | N-linked glycans | 23 | 11 | 11 | Steichen, 2019 |
| V2 apex | V2-int | PCT64-18D | HCDR3 | Loop binders | 1 | N-linked glycans | 25 | 17 | 8 | Landais, 2017 |

F

| Antibody | Paratope | $gp120_A$ | $gp120_B$ | $gp120_C$ | Glycans | Total |
|---|---|---|---|---|---|---|
| CD4bs-int | Heavy chain (Å) | 1014 | 90 | - | 140 | 1245 |
| | $V_H$-gene (%) | 85% | 100% | - | 100% | 87% |
| | HCDR3 (%) | 15% | 0% | - | 0% | 13% |
| | Light chain (Å) | 200 | 0 | - | 375 | 575 |
| | Total | 1214 | 90 | 0 | 516 | 1819 |
| V3-int | Heavy chain (Å) | 253 | - | - | 894 | 1147 |
| | $V_H$-gene (%) | 0% | - | - | 52% | 40% |
| | HCDR3 (%) | 100% | - | - | 48% | 60% |
| | Light chain (Å) | 507 | - | - | 236 | 743 |
| | Total (Å) | 760 | 0 | 0 | 1130 | 1890 |
| V2-int | Heavy chain (Å) | 371 | 346 | 239 | 817 | 1773 |
| | $V_H$-gene (%) | 8% | 0% | 0% | 60% | 29% |
| | HCDR3 (%) | 92% | 100% | 100% | 40% | 71% |
| | Light chain (Å) | 0 | 0 | 0 | 383 | 383 |
| | Total (Å) | 371 | 346 | 239 | 1200 | 2156 |

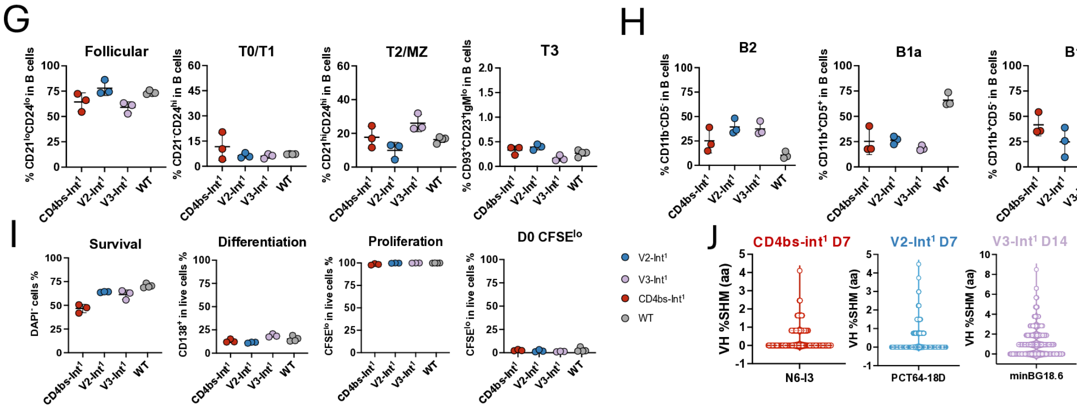


**Figure S1. Generation of KI mouse models for BCRs with distinct specificities against HIV Env, related to Figure 1.**

(A) Structural panel shows a representative CD4bs-int Fab (PDB:5FYJ, red) bound to the CD4bs epitope, with an inset depicting the $V_H$-dominant binding mode and the extensive contacts with the proteinaceous surface of gp120 Env.

(B) Structural panel shows a representative V2-int Fab (PDB:7T74, blue) bound to the V2 apex epitope, with an inset depicting the HCDR3-dominant binding mode and pronounced contribution of glycan contacts to the overall binding.

(C) Structural panel shows a representative V3-int Fab (PDB: 6DFG, purple) bound to the V3 glycan (N332) supersite, with an inset depicting the HCDR3-dominant binding mode and pronounced contribution of glycan contacts to the overall binding.

(D) Structural illustration of CD4bs-int (PDB:5FYJ, red), V3-int (PDB:6DFG, purple), and V2-int (PDB:7T74, blue) Fabs engaging the HIV Env trimer (PDB:7T74, light and dark grey) at their respective supersites, demonstrating that no steric clashes are expected upon simultaneous binding.

(E) Summary table comparing the binding modes of "loop binders" (V3-int, V2-int) and "$V_H$-binders" (CD4bs-int). V3-int and V2-int BCRs use HCDR3-dominant interactions with substantial N-linked glycan contact contributions, while CD4bs-int BCRs employ $V_H$-gene contacts to form extensive binding footprints on gp120. Blue gradient intensity represents relative buried surface area (BSA) at individual gp120 protomers (A, B, and C; grey) and N-linked glycans (beige). Green gradient intensity shows percent contribution to the heavy chain paratope by $V_H$-gene vs HCDR3 regions. Red gradient represents the relative paratope BSA distribution between gp120 residues and N-linked glycans (white = no contribution, dark color = highest contribution).

(F) Summary of HIV-specific bnAbs used to generate knockin mouse models, showing dominant paratope region, antibody:Env (Ab:Env) binding ratio, epitope specificity, HCDR3 length, and percent SHM at $V_H$ and $V_K$/$V_L$.

(G) Spleens from CD4bs-Int[1], V2-Int[1], V3-Int[1], and WT mice were analyzed by flow cytometry. B cells were divided into follicular B cells ($CD21^{lo}CD24^{lo}$), T0/T1 B cells ($CD21^{-}CD24^{hi}$), T2/MZ

B cells ($CD21^{hi}CD24^{hi}$), and T3 B cells ($CD93^{+}CD23^{+}IgM^{lo}$) and quantified for dot plots. (n = 3–4 mice per independent group). Error bars indicate mean ± SD from mice in pooled groups.

(H) Peritoneal lavage was performed on CD4bs-Int[1], V2-Int[1], V3-Int[1], and WT mice, and the exudate was analyzed by flow cytometry. B cells ($IgM^{+}$) were divided into (left) B2 ($CD11b^{-}CD5^{-}$), (middle) B1a ($CD11b^{+}CD5^{+}$), (right) B1b ($CD11b^{+}CD5^{-}$), and quantified for dot plots. (n = 3–4 mice per independent group). Error bars indicate mean ± SD from mice in pooled groups.

(I) Splenic B cells isolated from CD4bs-Int[1], V2-Int[1], V3-Int[1] as well as WT mice were individually purified and CFSE-labeled at day 0 and cultured and stimulated with CD40L, IL-4 and IL-5 in vitro. After 72 h, cultured B cells were analyzed by flow cytometry for (left) survival by percentage of live $DAPI^{-}$ cells, (middle) differentiation by percentage of $CD138^{+}$ cells, and (right) proliferation by percentage of $CFSE^{lo}$ cells (CFSE initial gate placed relative to day 0 for each line). Data are pooled from 3–4 biological replicates per group; Error bars indicate mean ± SD.

(J) Dotted violin plots of HC amino acid mutations in CD4bs-Int[1], V2-Int[1], or V3-Int[1] B cells at day 7 or 14 after immunization. Each dot represents an HC sequence from one B cell.

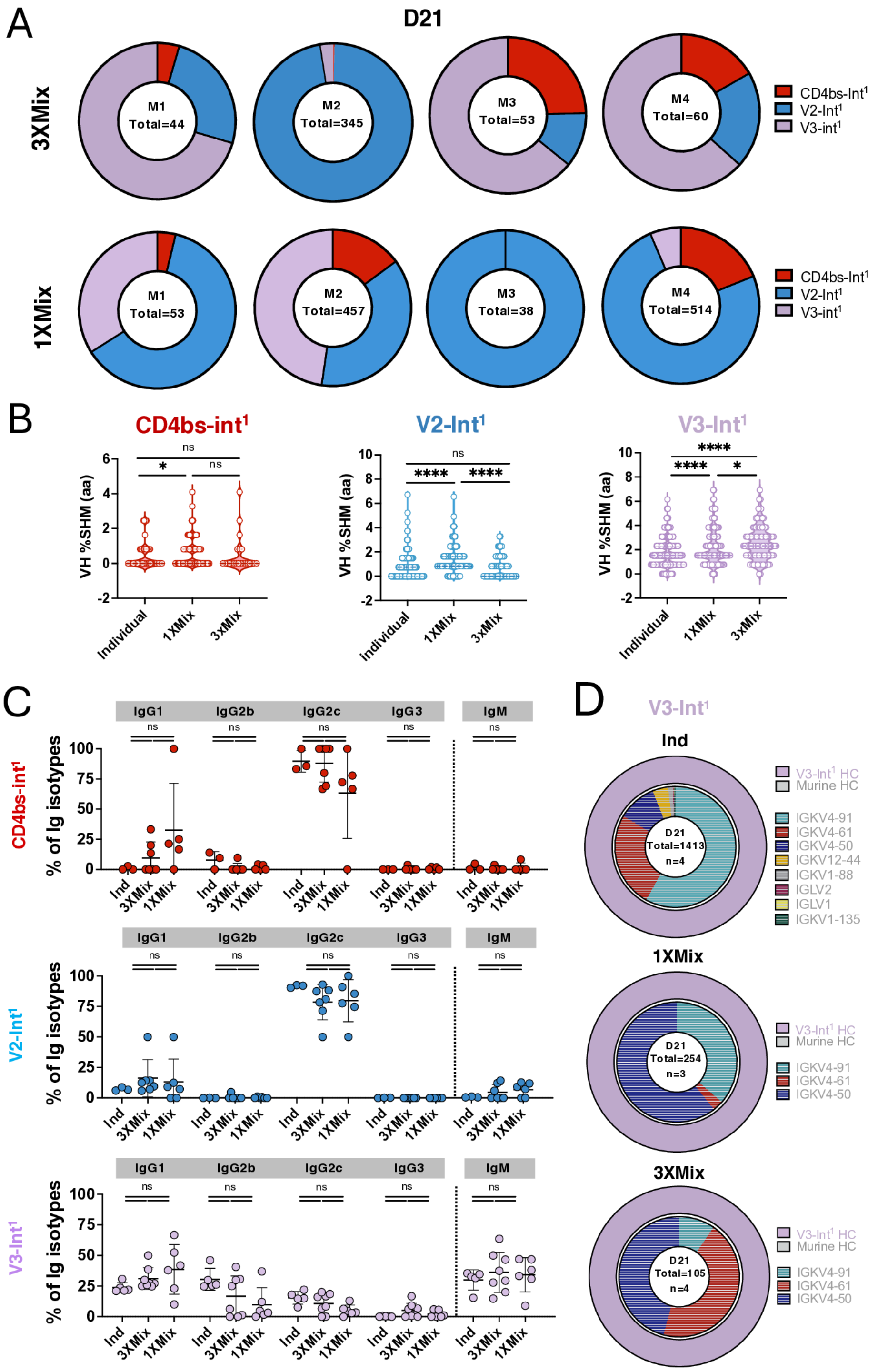


**Figure S2. Sequence analysis of three simultaneous responses targeting distinct epitopes, related to Figure 2.**

(A) B cell lineage frequency analysis from 10x scRNAseq of GC CD45.2 B cells sorted at day 21 from the 3x Mix (top) and 1x Mix (bottom) groups showing fraction of CD4bs-Int$^{1}$ (red), V2-Int$^{1}$ (blue), and V3-Int$^{1}$ (purple) sequences. Sequences are reproduced from **Fig. 2E** but broken out by individual mouse (Total=number of sequences amplified).

(B) Dotted violin plots of HC amino acid mutations in CD4bs-Int$^{1}$, V2-Int$^{1}$, or V3-Int$^{1}$ B cells in mixed transfer groups at day 21. Each dot represents HC sequence from one B cell. Sequence frequencies after individual transfer and immunization are reproduced from **Fig. 1H** for ease of direct comparison.

(C) Detailed class switch profiles at day 21 in individual (Ind) or Mix transfer recipients in (top) CD4bs-Int$^{1}$, (middle) V2-Int$^{1}$, and (bottom) V3-Int$^{1}$ recipient mice calculated from sequence data pulled from two individual experiments. Total sequences for CD4bs-Int$^{1}$: Ind=58, 3XMix=77, 1XMix=118; total sequences for V2-Int$^{1}$: Ind=892, 3XMix=231, 1XMix=415; total sequences for V3-Int$^{1}$: Ind=933, 3XMix=857, 1XMix=404.

(D) Nested pie chart showing V3-Int$^{1}$ LC usage from single-cell sorted epitope-specific (BG505$^{+}$KO$^{-}$) B cells at day 21 in mixed adoptive transfer recipients. Outer layer, human V3-Int$^{1}$ IGHV; inner layer, murine IGKV (n=number of sequences amplified).

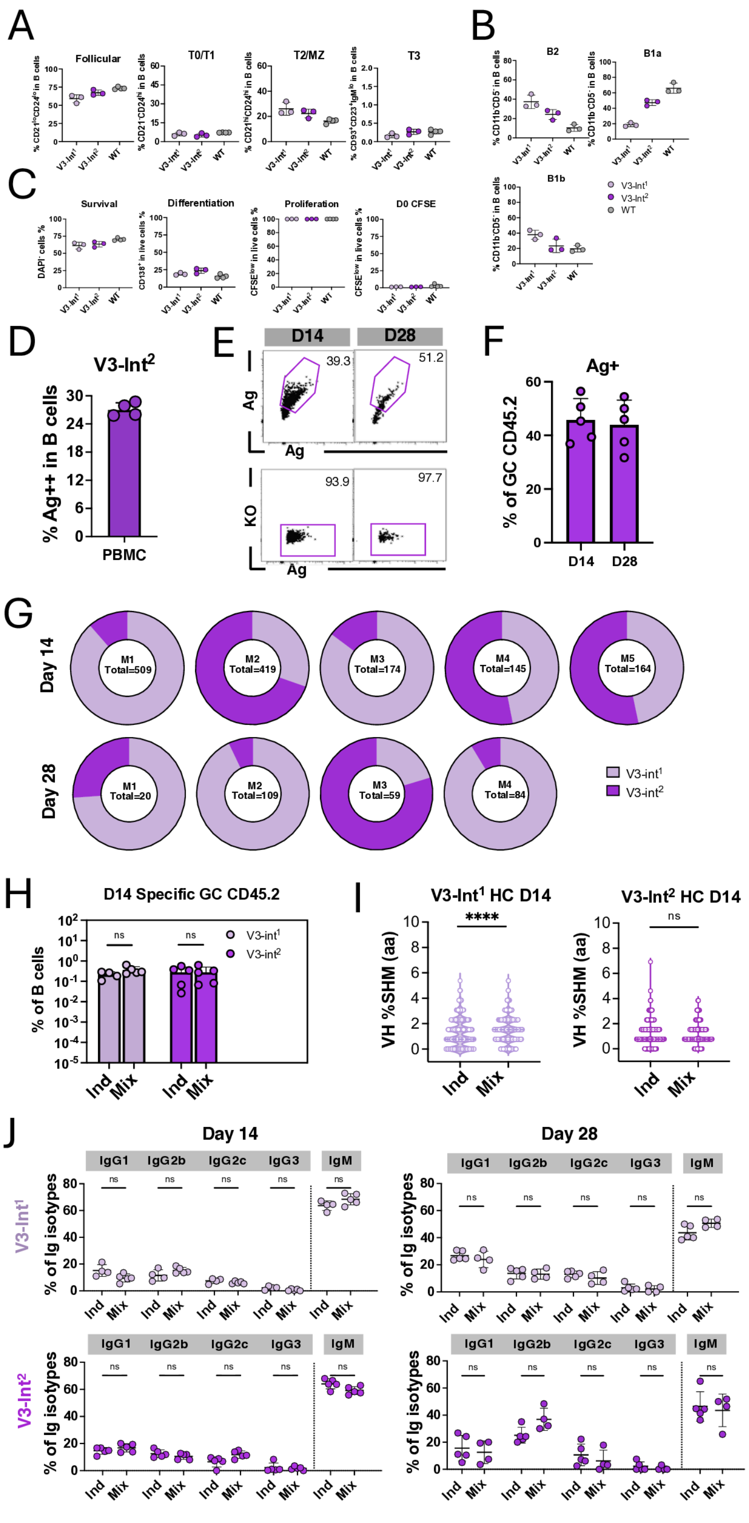


**Figure S3. Responses by low affinity cells targeting the same V3 epitope, related to Figure 3.**

(A) Spleens from V3-Int$^2$ mice were analyzed by flow cytometry. B cells were follicular B cells (CD21$^{lo}$CD24$^{lo}$), T0/T1 B cells (CD21$^-$CD24$^{hi}$), T2/MZ B cells (CD21$^{hi}$CD24$^{hi}$), and T3 B cells (CD93$^+$CD23$^+$IgM$^{lo}$) and quantified for dot plots (n = 3–4 mice per independent group). Error bars indicate mean ± SD from mice in pooled groups. Data for V3-Int$^1$ and WT group were reproduced from **Fig. S1G** for direct comparisons.

(B) Peritoneal lavage was performed on V3-Int$^2$ mice, and the exudate was analyzed by flow cytometry. B cells (IgM$^+$) were divided into (left) B2 (CD11b$^-$CD5$^-$), (middle) B1a (CD11b$^+$CD5$^+$), (right) B1b (CD11b$^+$CD5$^-$), and quantified for dot plots (n = 3–4 mice per independent group). Error bars indicate mean ± SD from mice in pooled groups. Data for V3-Int$^1$ and WT group were reproduced from **Fig. S1H** for direct comparisons.

(C) Splenic B cells isolated from V3-Int$^2$ mice were individually purified and CFSE-labeled at day 0 and cultured and stimulated with CD40L, IL-4 and IL-5 in vitro. After 72 h, cultured B cells were analyzed by flow cytometry for (left) survival by percentage of live DAPI$^–$ cells, (middle) differentiation by percentage of CD138$^+$ cells, and (right) proliferation by percentage of CFSE$^{lo}$ cells (CFSE initial gate placed relative to day 0 for each line). Data are pooled from 3–4 biological replicates per group; Error bars indicate mean ± SD. Data for V3-Int$^1$ and WT group were reproduced from **Fig. S1I** for direct comparisons.

(D) BG505-specific peripheral blood B cells in V3-Int$^2$ KI mice (n = 4). Dots represent individual animals and bars are mean + SD.

(E) Representative FACS plots of B cells obtained from dLNs of V3-Int$^2$-transferred mice at 14 and day 28 with BG505 trimer. (upper) Events pre-gated on lymphocytes/singlets/live/CD4$^-$CD8$^-$F4/80$^-$Gr1$^-$/B220$^+$/CD38$^-$CD95$^+$/CD45.2$^+$CD45.1$^-$ B cells and represent Ag binders in GC CD45.2.

(lower) Events pre-gated on lymphocytes/singlets/live/$CD4^{-}CD8^{-}F4/80^{-}Gr1^{-}/B220^{+}/CD38^{-}$ $CD95^{+}/CD45.2^{+}CD45.1^{-}/ Ag^{++}$ B cells and represent BG505-V3-$KO^{-}$ cells in GC CD45.2 binders.

(F) BG505 probe binders in GC CD45.2 cells. Dots represent individual animals and bars are mean + SD.

(G) B cell lineage frequency analysis from 10x scRNAseq of GC CD45.2 B cells sorted from mix-transfer recipients at days 14 and 28 showing fraction of V3-$Int^1$ (light purple) and V3-$Int^2$ (dark purple) sequences. Sequences are reproduced from **Fig. 3I** but broken out by individual mouse (Total=number of sequences amplified).

(H) Specific GC CD45.2 B cells as percentage in total B cells. Percentage was calculated from FACS data for individual groups, and from 10x scRNAseq plus FACS data for Mix-transfer groups. Values of zero were plotted as UD on the $log_{10}$ scale. Adjusted p-values (q-values) calculated by Kruskal Wallis test with pairwise post-hoc comparisons adjusted using BYK correction. ns > 0.05.

(I) Dotted violin plot of HC amino acid mutations across all sites at days 14 and 28. Each dot represents an HC sequence from one B cell. Statistical analysis was performed using Mann-Whitney's test. P-values are indicated as follows: ****$p < 0.0001$; ns = not significant.

(J) Detailed class switch profiles of (top) V3-$Int^1$ and (bottom) V3-$Int^2$ at days 14 and 28 in individual or mix transfer recipients. Total sequences for D14 V3-$Int^1$: Ind=609, Mix=693; total sequences for D14 V3-$Int^2$: Ind=912, Mix=671; total sequences for D28 V3-$Int^1$: Ind=471, Mix=177; total sequences for D28 V3-$Int^2$: Ind=237, Mix=96.

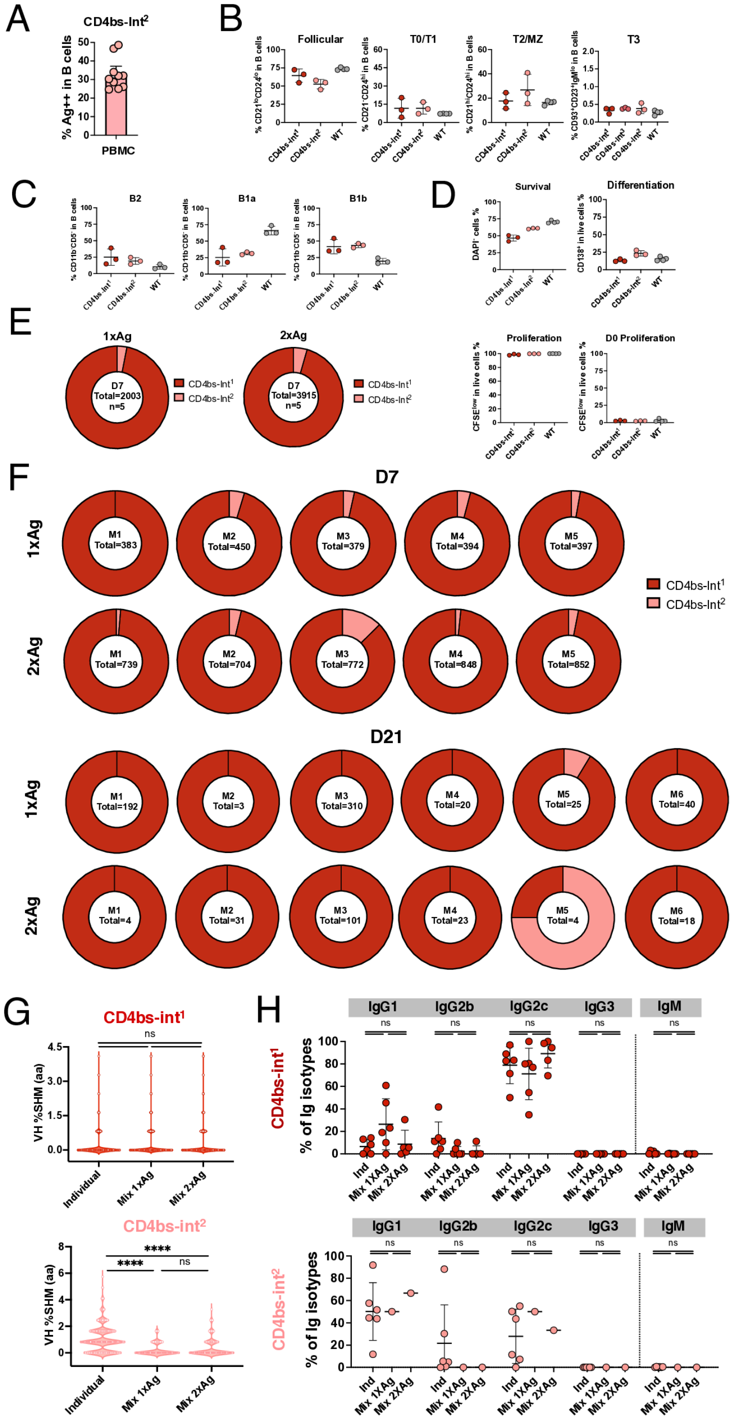


**Figure S4. Response to immunization in CD4bs-Int$^1$ and CD4bs-Int$^2$ mix-transfer recipients compared to individual transfer recipients, related to Figure 4.**

(A) BG505-specific peripheral blood B cells in CD4bs-Int$^{2}$ KI mice. Dots represent individual animals and bars are mean + SD. Ten animals from four different experiments were pooled for analysis.

(B) Spleens from CD4bs-Int$^{2}$ mice were analyzed by flow cytometry. B cells were divided into follicular B cells (CD21$^{lo}$CD24$^{lo}$), T0/T1 B cells (CD21$^{-}$CD24$^{hi}$), T2/MZ B cells (CD21$^{hi}$CD24$^{hi}$), and T3 B cells (CD93$^{+}$CD23$^{+}$IgM$^{lo}$) and quantified for dot plots (n = 3–4 mice per independent group). Error bars indicate mean ± SD from mice in pooled groups. Data for CD4bs-Int$^{1}$ and WT group were reproduced from **Fig. S1G** for direct comparisons.

(C) Peritoneal lavage was performed on CD4bs-Int$^{2}$ mice, and the exudate was analyzed by flow cytometry. B cells (IgM$^{+}$) were divided into (left) B2 (CD11b$^{-}$CD5$^{-}$), (middle) B1a (CD11b$^{+}$CD5$^{+}$), (right) B1b (CD11b$^{+}$CD5$^{-}$), and quantified for dot plots (n = 3–4 mice per independent group). Error bars indicate mean ± SD from mice in pooled groups. Data for CD4bs-Int$^{1}$ and WT group were reproduced from **Fig. S1H** for direct comparisons.

(D) Splenic B cells isolated from CD4bs-Int$^{2}$ mice were individually purified and CFSE-labeled at day 0 and cultured and stimulated with CD40L, IL-4 and IL-5 in vitro. After 72 h, cultured B cells were analyzed by flow cytometry for (left) survival by percentage of live DAPI$^{–}$ cells, (middle) differentiation by percentage of CD138$^{+}$ cells, and (right) proliferation by percentage of CFSE$^{lo}$ cells (CFSE initial gate placed relative to day 0 for each line). Data are pooled from 3–4 biological replicates per group; error bars indicate mean ± SD. Data for CD4bs-Int$^{1}$ and WT group were reproduced from **Fig. S1I** for direct comparisons.

(E) The CD4bs-Int$^{2}$ (pink) and CD4bs-Int$^{1}$ (red) and CD4bs-Int$^{2}$ (pink)B cell lineage frequency analysis from 10x scRNAseq of GC CD45.2 B cells sorted from mix-transfer recipients at day 7 after 1X or 2X Ag dose. Pie charts are averages of all mice in a group (Total=number of sequences amplified, n=number of mice).

(F) B cell lineage frequency analysis from 10x scRNAseq of GC CD45.2 B cells sorted from mix-transfer recipients at days 7 and 21 after 1X or 2X Ag dose showing fraction of CD4bs-Int$^{1}$ (red) and CD4bs-Int$^{2}$ (pink) sequences. Sequences are reproduced from **Fig. 4H** but broken out by individual mouse (Total=number of sequences amplified).

(G) Dotted violin plot of HC amino acid mutations at day 21. Each dot represents an HC sequence from one B cell. Statistical analysis performed using Kruskal Wallis test with pairwise post-hoc comparisons adjusted using BYK correction. Adjusted p-values (q-values) are indicated as follows: ****$p < 0.0001$; ns =>0.05.

(H) Detailed class switch profiles at day 21 in individual or mix transfer recipients in (top) CD4bs-Int$^{1}$ and (bottom) CD4bs-Int$^{2}$ recipient mice. Total sequences for CD4bs-Int$^{1}$: Ind=315, Mix1XAg=318, Mix 2XAg=200; total sequences for CD4bs-Int$^{2}$: Ind=1398, Mix1XAg=2, Mix 2XAg=3.

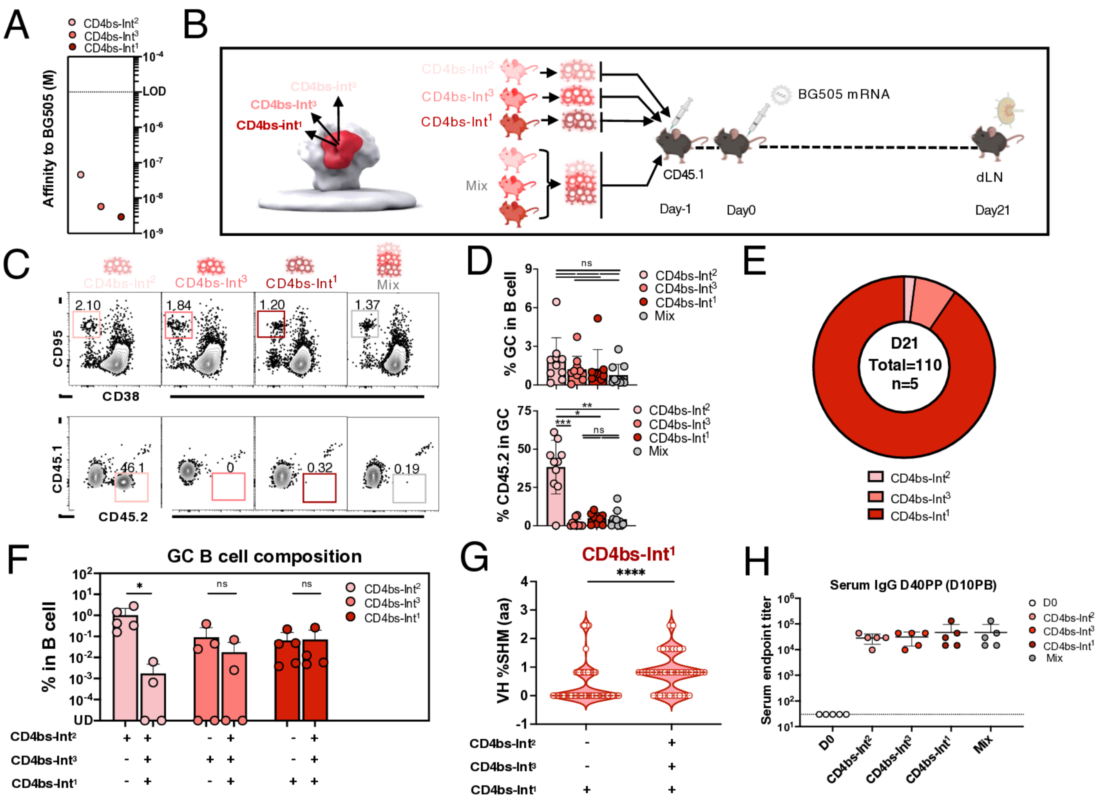


**Figure S5. Response targeting the same CD4bs epitope by three B cell lines of variable affinities, related to Figure 4.**

(A) Affinity of the CD4bs-Int$^1$, CD4bs-Int$^2$, and CD4bs-Int$^3$ against BG505 trimer measured by SPR dissociation constant. Each dot represents the mean of four technical replicates. Dotted line represents LOD. Affinities of CD4bs-Int$^1$ and CD4bs-Int$^2$ reproduced from **Figs. 1D and 4A** for direct comparison.

(B) Schematic of individual or mixed-adoptive-transfer and immunization experiments for three lines targeting the same CD4bs epitope.

(C) Representative FACS plots of B cells obtained from dLNs at 21 days post-immunization by BG505 in mice transferred with different cells. Events were pre-gated on lymphocytes/singlets/live/CD4$^-$CD8$^-$F4/80$^-$Gr1$^-$/B220$^+$ B cells and represent GC in B cells or CD45.2 cells in GC.

(D) (upper) GC cells as the percentage of total B cells and (lower) $CD45.2^+$ KI cells as the percentage of total GC B cells at day 21. Dots represent individual values pooled from two experiments and bars are mean ± SD. Adjusted p-values (q-values) calculated by Kruskal Wallis test with pairwise post-hoc comparisons adjusted using BYK correction. Adjusted p-values (q-values) are indicated as follows: *p < 0.05, ns = not significant.

(E) The CD4bs-$Int^3$ (rose), CD4bs-$Int^2$ (pink), and CD4bs-$Int^1$ (red) B cell lineage frequency analysis from 10x scRNAseq of GC CD45.2 B cells sorted at day 21 from Mix-transfer recipients. Pie charts are averages of all mice in a group.

(F) BCR composition of CD45.2 GC B cells as a percentage of total B cells. Percentage was calculated from FACS data for individual groups, and from 10x scRNAseq plus FACS data for Mix-transfer recipients. Dots represent individual values and bars are mean ± SD. Values of zero were plotted as UD on the $\log_{10}$ scale. Adjusted p-values (q-values) calculated by 2-way ANOVA test with pairwise post-hoc comparisons adjusted using BYK correction. Adjusted p-values (q-values) are indicated as follows: *p < 0.05, ns = not significant.

(G) Dotted violin plot of HC amino acid mutations across all sites 21 days post-immunization by BG505 mRNA. Each dot represents HC sequence from one B cell. Sequence frequencies after individual transfer and immunization are reproduced from **Fig. 1H** for ease of direct comparison.

Statistical analysis made using Mann-Whitney's test. P-values are indicated as follows: ****p < 0.0001

(H) ELISA quantification of BG505-binding IgG from individual or Mix transfer-recipient mice immunized with BG505 mRNA and then boosted with 1 μg of a BG505 mRNA homologous boost at day 30. Samples were taken pre-immunization (day 0) or ten days post-boost. Dots represent individual animals and bars are mean ± SD.

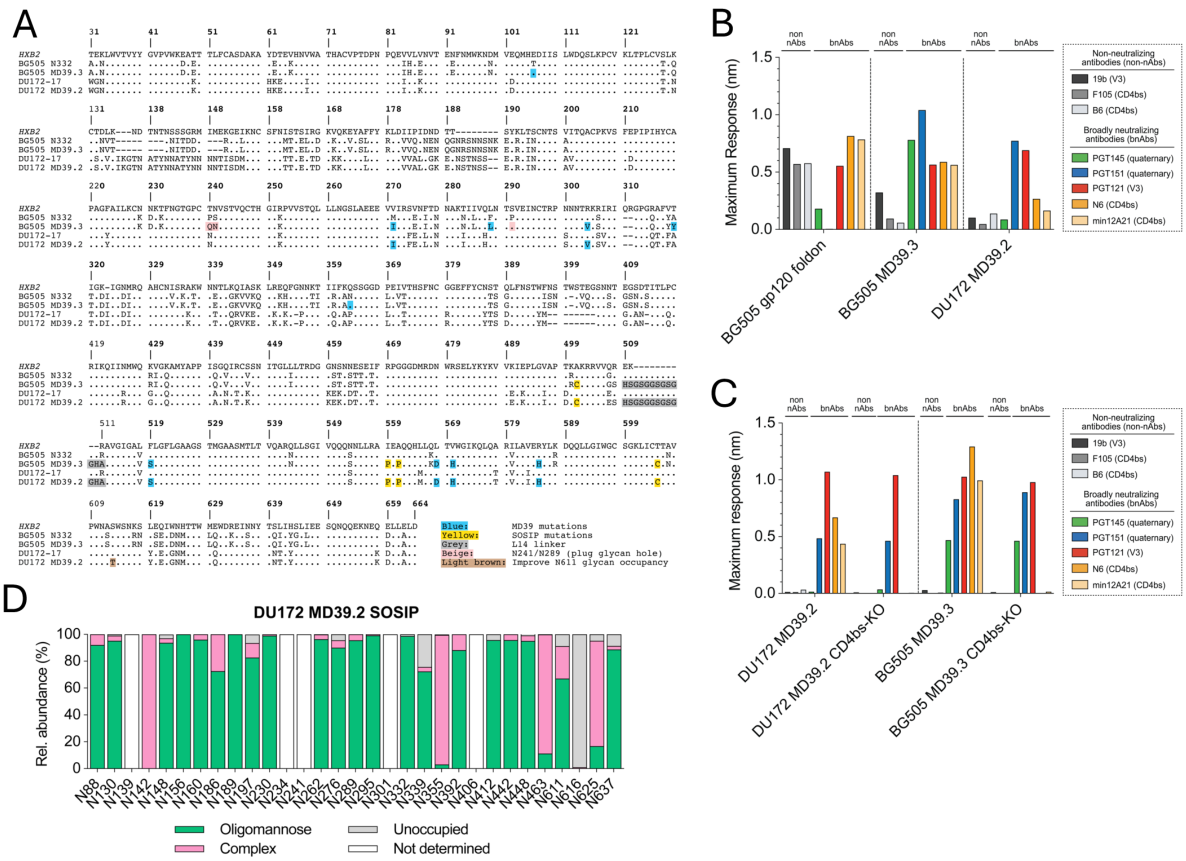


**Figure S6. Characterization of DU172 immunogen and probe, related to Figure 5 and Methods.**

(A) Sequence alignment of stabilized gp140 immunogens (BG505 MD39.3 and DU172 MD39.2) and corresponding wild-type HIV-1 isolates aligned to HxB2 reference sequence, with color-coded MD39 (blue), SOSIP (yellow), glycan masking (beige), link14 linker (grey), and glycan occupancy-enhancing (light brown) mutation.

(B) Antigenic profile of DU172 MD39.2 by BLI using non-neutralizing antibodies (non nAbs) (CD4bs-specific F105 and B6; V3-specific 19b) and bnAbs (quaternary-specific PGT151 and PGT145; V3-specific PGT121; CD4bs-specific N6 and min12A21) confirmed native-like conformation of the stabilized gp140 trimer. Non nAbs preferentially bind open Env conformations; BG505 gp120 foldon served as a negative control representing poorly assembled trimers. Since PGT145 does not neutralize DU172-17, it is therefore not a valid positive control for assessing DU172 trimer conformation.

(C) Bio-layer interferometry (BLI) was used to assess antigenic profiles of CD4bs-WT and VRC01-class KO sort reagents. CD4bs-specific knockout mutations in DU172 MD39.2 and BG505 MD39.3 effectively block VRC01-class bnAb recognition. Binding to quaternary structure-dependent antibodies (PGT151, PGT145) and lack of binding to non-neutralizing antibodies (V3-specific 19b; CD4bs-specific F105 and B6) confirmed proper folding.

(D) Site-specific N-glycan processing of stabilized DU172 MD39.2 immunogen reveals high glycan occupancy at all PNGS sites except N616, with the S613T mutation effectively improving N611 glycan occupancy. Relative abundance is shown as percentage of high mannose (green), complex (pink), unoccupied (grey), and not determined (white) at all glycan sites.

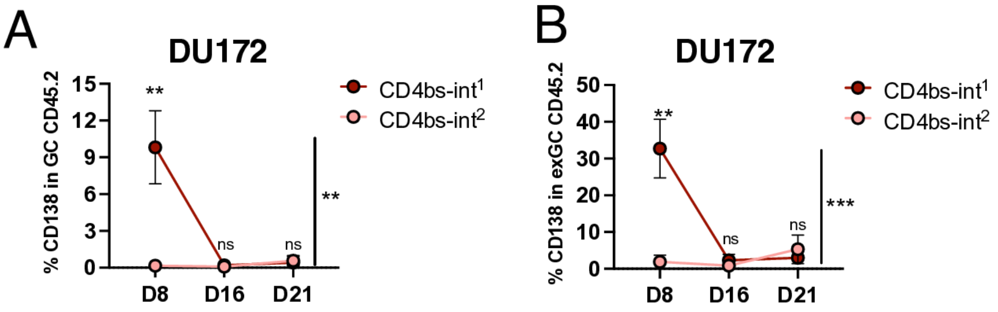


**Figure S7. Response of CD4bs-Int$^1$ and CD4bs-Int$^2$ after DU172 immunization, related to Figure 7.**

(A–B) Time course plots of CD138$^+$ plasma cells as a percentage of total (A) in GC CD45.2$^+$ cells or (B) exGC CD45.2$^+$ cells. Data obtained from dLNs in mice transferred with CD4bs-Int$^1$ or CD4bs-Int$^2$ 8, 16, and 21 days post-immunization by DU172 protein adjuvanted with SMNP. Dots are average and bars are mean ± SD. Adjusted p-values (q-values) calculated by 2-way ANOVA test with pairwise post-hoc comparisons adjusted using BYK correction. Adjusted p-values (q-values) are indicated as follows: *$p < 0.05$, **$p < 0.01$, ***$p < 0.001$, ****$p < 0.0001$; ns = not significant.

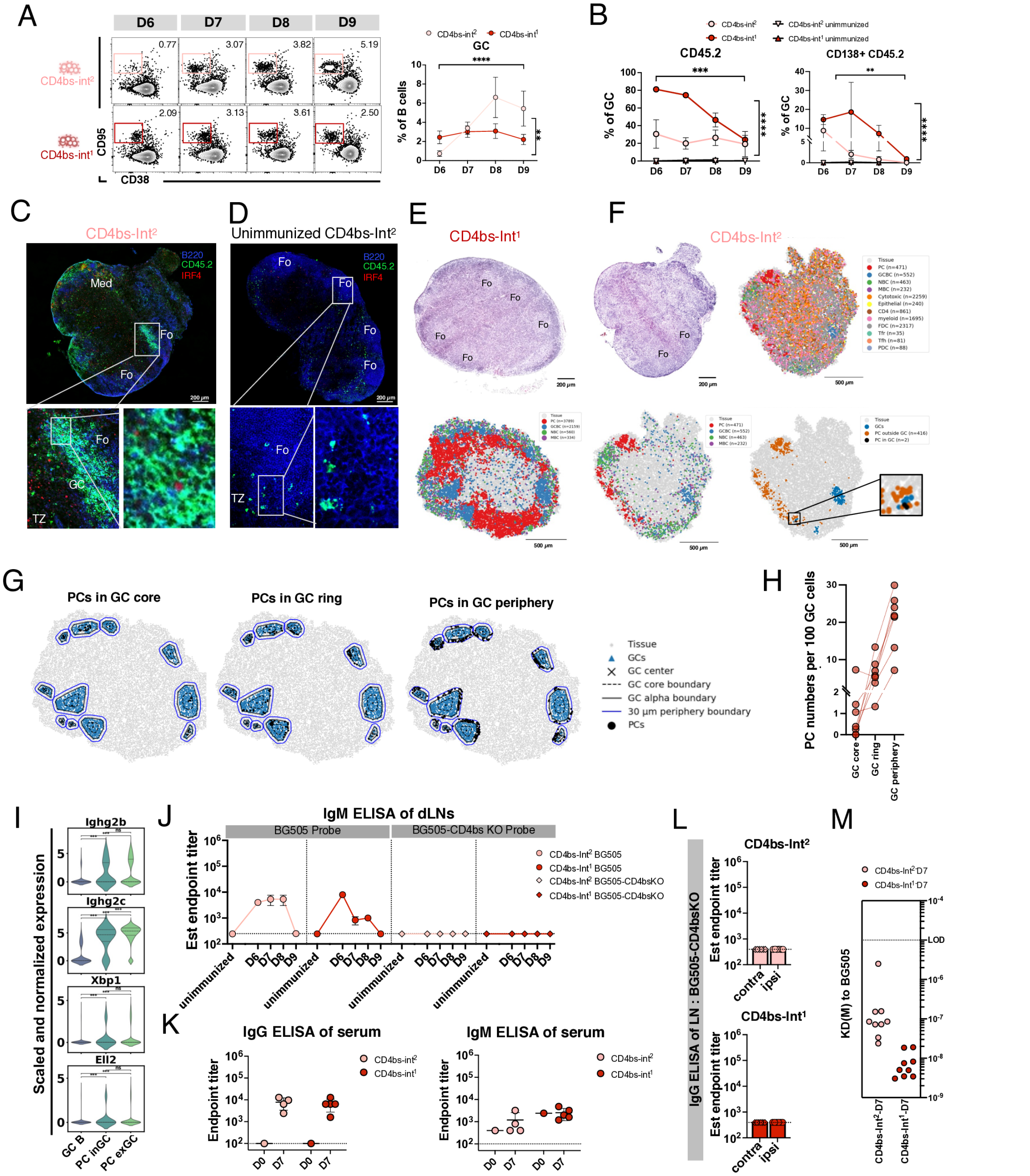

**Figure S8. Characterization of PC-like cells in dLN in CD4bs-Int¹ or CD4bs-Int² recipient mice, related to Figure 7.**

(A–B) Data obtained from dLNs in mice transferred with CD4bs-Int$^{1}$ or CD4bs-Int$^{2}$ at 6–9 days post-immunization by BG505 mRNA. Dots are means and bars are mean ± SDs. P-values calculated by 2-way ANOVA test.

(A) (left) Representative FACS plots of GC in B cells. Events were pre-gated on lymphocytes/singlets/live/CD4$^{-}$CD8$^{-}$F4/80$^{-}$Gr1$^{-}$/B220$^{+}$ B cells and represent GC in B cells. (right) Time course plots of GC cells as percentage of total B cells.

(B) Time course plots of (left) CD45.2 cells as percentage of GC B cells and (right) CD138+ CD45.2 B cells as percentage of GC B cells.

(C) Representative immunofluorescent staining of dLN isolated from mice with CD4bs-Int$^{2}$ B cells 8 days post-immunization by BG505, staining with B220 (blue, surface), CD45.2 (green, surface), and IRF4 (red, nucleus). Large tile presents overview of architecture of the whole LN, inset boxes present individual GCs or subsets of PCs in GC. Fo, Follicle; MC, Medullar cords; TZ, T cell zone. Scale bar: 200 µm.

(D) Representative immunofluorescent staining of dLN isolated from unimmunized control mice transferred with CD4bs-Int$^{2}$ B cells at day 8, staining with B220 (blue, surface), CD45.2 (green, surface), and IRF4 (red, nucleus). Large tile presents overview of architecture of the whole LN, inset boxes present individual Follicle or subsets of B cells in the Follicle. Fo, Follicle; TZ, T cell zone. Scale bar: 200 µm.

(E) (upper) Representative hematoxylin and eosin (H&E) staining section of dLN isolated from mice transferred with CD4bs-Int$^{1}$ B cells 6 days post-immunization by BG505. Fo, Follicle. Scale bar: 200 µm. (lower) Spatial map of dLN B cells colored by subtype. Scale bar: 500 µm. n= number of subset cells. Non-serial sections of the same dLN were used for fluorescent-staining, H&E staining and Slide-seq assay.

(F) (upper left) Representative H&E staining section of dLN isolated from mice transferred with CD4bs-Int[2] B cells 8 days post-immunization by BG505. Scale bar: 200 μm. Spatial map of dLN isolated from mice with CD4bs-Int[1] B cells at 6 days post-immunization BG505 by Slide-seq, colored by (lower left) B cell subtypes, (upper right) all cell types, or (lower right) PC anatomical location. Scale bar: 500 μm. PC, Plasma cells; GCBC, GC B cells; NBC, naïve B cells; MBC, memory B cells; Cytotoxic, cytotoxic T cells; Epithelial, epithelial cell; CD4, CD4 T cells; FDC, follicular dendritic cells; Tfr, follicular regulatory T cells; Tfh, follicular helper T cells; PDC, plasmacytoid dendritic cells. n=number of subset cells. Non-serial sections of the same dLN were used for fluorescent-staining, H&E staining and Slide-seq assay.

(G) PC spatial distribution in (left) GC core, (center) GC ring and (right) GC periphery from dLN isolated from mice transferred with CD4bs-Int[1] B cells 6 days post-immunization by BG505. The inner "GC core" region was created by homothetically shrinking the GC boundary to half of its original size about the center. the outer "GC ring" region was defined by the remaining area between the inner GC core region and the GC outer boundary. The "GC periphery" region was created by expanding a 30 μm boundary around each GC using Minkowski addition. Each black dot represents a single PC; blue triangles represent GC cells.

(H) Quantification of PC spatial distribution in GC core, GC ring and GC periphery from (G). The number of PCs at each site were scaled to the size (GC cell number) of the same GC of sufficient size (>= 70 beads), to get the PC numbers per 100 GC cells for each GC. Dots represent GCs, lines link numbers from the same GC.

(I) Violin plot showing selected gene expression in GC B cells, PCs inGC, and PCs exGC. Pairwise group comparisons were performed using two-sided Mann–Whitney U tests, followed by Holm–Bonferroni correction for multiple comparisons.

(J) ELISA quantification of BG505-binding (circle) and BG505-CD4bs-KO-binding (diamond) IgM from 100 µl of dLN homogenates of CD4bs-In[1] (red) or CD4bs-Int[2] (pink) recipient mice immunized with BG505 mRNA or left unimmunized. A 5x dilution multiplier was applied to obtain the final est endpoint titer of dLNs. Dots represent mean values of triplicate technical replicates from homogenates generated from one popliteal LN from each of the four mice pooled into 100 µl of buffer. Bars are mean ± SD. Unimmunized dots include all data collected at days 7, 8 and 9 in groups left unimmunized after corresponding adoptive transfers. Dotted line represents LOD.

(K) ELISA quantification of BG505-binding IgG (upper) and IgM (lower) from serum of CD4bs-In[1] (red) or CD4bs-Int[2] (pink) recipient mice either pre-immunization (day 0) or 7 days post-immunization by BG505. Dots represent individual animals and bars are mean ± SD. Dotted line represents LOD.

(L) ELISA quantification of BG505-CD4bsKO-binding IgG from 100 µl of dLN homogenates of ipsilateral dLNs (circle) and contralateral LNs (diamond) of CD4bs-In[1](red) or CD4bs-Int[2] (pink) recipient mice immunized with BG505 mRNA or left unimmunized. A 5x dilution multiplier was applied to obtain the final est endpoint titer of dLNs. For each group, homogenates were generated from one popliteal LN from each of four mice pooled into 100 µl of buffer. Technical triplicates were performed for each group. Dots represent individual values pooled from two experiments. Bars are mean ± SD. Dotted line represents LOD.

(M) SPR affinity against BG505 for Abs sorted from D7 GC for CD4bs-In[1](red) or CD4bs-Int[2] (pink) lineages in individual transfer experiments from **Fig. 4E**. Each dot represents the mean of 4 technical replicates. Dotted line represents LOD.

Adjusted p-values (q-values) are indicated as follows: $^{**}p < 0.01$, $^{***}p < 0.001$; ns = not significant.
